# Id2 levels determine the development of effector vs. exhausted tissue-resident memory CD8^+^ T cells during CNS chronic infection

**DOI:** 10.1101/2024.07.30.605547

**Authors:** Aboubacar Sidiki K. Coulibaly, Lucie Nozeran, Céline Thomann, Marine Alis, Emilie Bassot, Ali Hassan, Rémi Porte, Marcy Belloy, Nicolas Blanchard, Frederick Masson

**Author notes:** 3d.FAB, CNRS, INSA, CPE-Lyon, UMR5246, ICBMS, University Claude Bernard Lyon 1, Villeurbanne, France.

## Abstract

Tissue-resident memory T cells (Trm) are essential for regional immunity in non-lymphoid tissues. Although single-cell transcriptomics have revealed Trm heterogeneity in various diseases, the molecular mechanisms behind this diversity are unclear. To investigate this, we used *Toxoplasma gondii* (*T. gondii*) infection, which persists in the central nervous system (CNS) and is controlled by brain CD8^+^ Trm. Our single-cell transcriptomic analysis of brain CD8^+^ T cells from *T. gondii*-infected mice showed heterogeneous expression of the transcriptional regulator Id2, correlating with different functional states. Using mixed bone marrow chimeras, we found that Id2-deficiency in T cells caused parasite-specific Trm to develop an altered phenotype with diminished effector functions and reduced expression of CD49a. Furthermore, loss of Id2 in brain-infiltrating CD8^+^ T cells led to the accumulation of exhausted PD1^+^Tox^+^CD8^+^ Trm cells, while Id2 overexpression repressed T cell exhaustion. Overall, our study shows that Id2 levels dictate the acquisition of effector *vs.* exhausted phenotypes in CD8^+^ Trm during chronic CNS infection.

**One sentence Summary:** Id2 expression level regulates the functional heterogeneity of brain Trm during CNS chronic infection

## INTRODUCTION

In recent years, tissue-resident memory T cells (Trm) have been identified as a crucial T cell population stably residing within non-lymphoid tissues and playing an essential role to protect against pathogen reinfections (*1, 2*). Initially recognized for their protective role against re-exposure to acute infections, Trm cells have also been found to exert important functions in situations of antigenic persistence (*3*). Indeed, these Trm-like cells play a protective role to control chronic latent pathogen infection (*4, 5*), and their presence has been correlated with good prognosis in solid cancers (*6*). Besides these protective functions, Trm can also play a deleterious role during chronic inflammatory and autoimmune diseases such as psoriasis, type I diabetes, or CNS autoimmune diseases (*7–9*).

Trm retention in non-lymphoid organs depends on the establishment of a tissue residency transcriptional program orchestrated by key transcriptional regulators including Blimp-1 and its homolog Hobit, as well as Runx3 and Bhlhe40 (*2*). The implementation of this tissue-residency program results in the induction of tissue retention associated molecules such as CD69, CXCR6, CD49a and CD103, along with the downregulation of tissue egress signals mediated by S1PR1 and KLF2, and a tissue-specific metabolic adaptation to the target tissue (*2*). Importantly, recent transcriptomic analysis have revealed the phenotypic and functional heterogeneity of Trm in various pathophysiological contexts (*5, 10–15*). Indeed, Trm subsets with distinct transcriptional signatures have been shown to coexist within a given tissue including “effector-like” Trm with increased effector potential, exhausted Trm exhibiting classical phenotypical features of T cell exhaustion and “stem-like” Trm potentially capable of self-replenishment (*5, 10, 11*). The heterogeneity of Trm is likely influenced by variations in cytokine signalling, the extent of CD4^+^ T cell help, and a differential access to specific antigen. However, the molecular mechanisms driving such phenotypical and functional diversity are still poorly understood.

In CD8^+^ T cells, these signals are typically integrated at the transcriptional level by a handful of key transcriptional regulators. Recent studies have shown that transcription factors important for effector CD8^+^ T cell differentiation such as T-bet, Blimp-1 or Runx3, also played an important role in regulating the development of Trm highlighting the intricate link between the effector and the Trm transcriptional programs of CD8^+^ T cell differentiation (*16–22*). We and others have previously demonstrated that the lineage fate decision into effector or memory CD8^+^ T cells during an acute and cleared pathogen infection is regulated by the interplay between Inhibitor of DNA binding 2 (Id2) transcriptional regulator and the class I basic helix-loop-helix protein (bHLH) family of transcription factors called E proteins, including E2A, HEB and E2-2 (*23–26*). Id2 is upregulated in effector CD8^+^ T cells during an acute viral infection and its loss cripples effector differentiation and programs CD8^+^ T cells to adopt a memory fate through the E2A-mediated activation of *Tcf7* (encoding Tcf1) (*24, 25*). While the functions of E2A and its inhibitor Id2 have been investigated during acute viral infection, much less is known about the role of this transcriptional axis in CD8^+^ Trm cells responding to a tissue-restricted chronic infection. Id2 has been shown to contribute to the homeostasis of intestinal Trm/intraepithelial lymphocytes (IEL) after an acute and resolved LCMV infection, and accordingly Id2 full knock-out mice lack IEL (*11, 27*). Whether and how Id2 plays a role in Trm biology in others peripheral tissues is unknown. The role of Id2 in T cell exhaustion is also still debated as its expression is increased in terminally exhausted cells during LCMV chronic infection and in terminally exhausted tumor-infiltrating lymphocytes (TILs) (*28, 29*). On the other hand, during chronic LCMV infection, Id2-deficiency promotes the differentiation of CD8^+^ T cells that show features of progenitors of exhausted T cells (Tpex) with increased Tcf1 expression (*30*). Finally, Tox the master regulator of the transcriptional and epigenetic remodelling of T cell exhaustion (*28, 31, 32*) has been shown to repress Id2 in a model of CNS autoimmunity (*33*), and accordingly Tox-forced expression decreased Id2 expression in progenitors of exhausted T cells (*28*). Collectively, these studies suggest a potential role of Id2 in regulating Trm biology and T cell exhaustion. In this study, we have explored the role of Id2 and its interplay with E2A in controlling Trm fate in a model of chronic *T.gondii* infection that establishes a latent infection in the CNS. Our data showed that Id2 is expressed in Trm and it repressed their acquisition of an exhaustion-associated phenotype, while promoting the expression of effector functions and tissue-retention molecules.

## RESULTS

### Id2 expression level correlates with the effector potential of brain CD8^+^ Trm during chronic *T. gondii* infection

To understand the role of Id2 in Trm biology during chronic *T.gondii* infection, we first wished to determine its expression pattern in CD8^+^ T cells responding to *T.gondii* infection. To this end, we performed a single-cell transcriptomic analysis (scRNA-seq) of CD8^+^ T cells isolated from the brain and the spleen of mice chronically infected with *Toxoplasma gondii*.GRA6-OVA (Tg.GRA6-OVA) (*34*) (**Fig. 1A**). We have recently shown using this infection model that following the acute phase of infection, *T. gondii* establishes a latent infection in the brain (from day 21 post-infection onward) that is efficiently controlled by parasite-specific brain CD8^+^ Trm (*5*). Unsupervised clustering analysis uncovered 6 CD8^+^ T cell clusters in the brain (cluster (cl)2, cl3, cl6, cl7, cl8 and cl9) and 4 clusters in the spleen (cl0, cl1, cl4 and cl5) (**Fig. 1B**). Notably, cl2, cl3, cl6 and cl7 had low expression of the tissue egress-associated molecules *Klf2* and *S1pr1,* while they expressed various levels of Trm-associated markers among which *Cd69*, *Cxcr6*, *Itga1* (encoding CD49a) or *Itgae* (encoding CD103) suggestive of a Trm phenotype (**Fig. 1C**). In line with this, gene-set enrichment analysis (GSEA) showed an enrichment of previously published core Trm and brain Trm gene signatures (*11, 35*) in cells from cl2, cl3, cl6, and cl7 (**Fig. 1D**). Aside from these 4 Trm clusters, which accounted for more than 94% of the brain-infiltrating CD8^+^ T cells, we identified a small Tcm cluster characterized by high *Tcf7* expression within the brain (cl8), and a cluster of terminal effector cells marked by high *Klrg1* expression (cl9), which was also found in the spleen (cl5) (**Fig. 1C)**. Notably, Id2 was expressed in all brain Trm clusters although its expression was heterogenous, with the highest expression observed in cl2 (**Fig. 1C).** Indeed, among the two most abundant Trm clusters, cl2 (42% of brain-infiltrating CD8^+^ T cells) and cl3 (39% of brain-infiltrating CD8^+^ T cells), we found that the expression of Id2 was significantly higher in cl2 compared with cl3 (**Fig. 1E**). Id2 was also highly expressed within terminal effector T cell clusters from the brain (cl9) and spleen (cl5) as expected from previous studies (*24, 25*). Since Id2 was differentially expressed between the two most abundant Trm clusters cl2 and cl3 (**Fig. 1E**), we asked whether this difference of expression may directly impact their transcriptome. GSEA comparison between cl2 and cl3 showed an enrichment of a previously published Id2-induced gene signature (*25*) in cl2, and its depletion in cl3. Conversely, Id2-repressed gene signature was positively enriched in cl3 and depleted in cl2 suggesting a direct impact of Id2 expression level in regulating the transcriptome of these Trm clusters (**Fig. 1F**).

**Figure 1:**
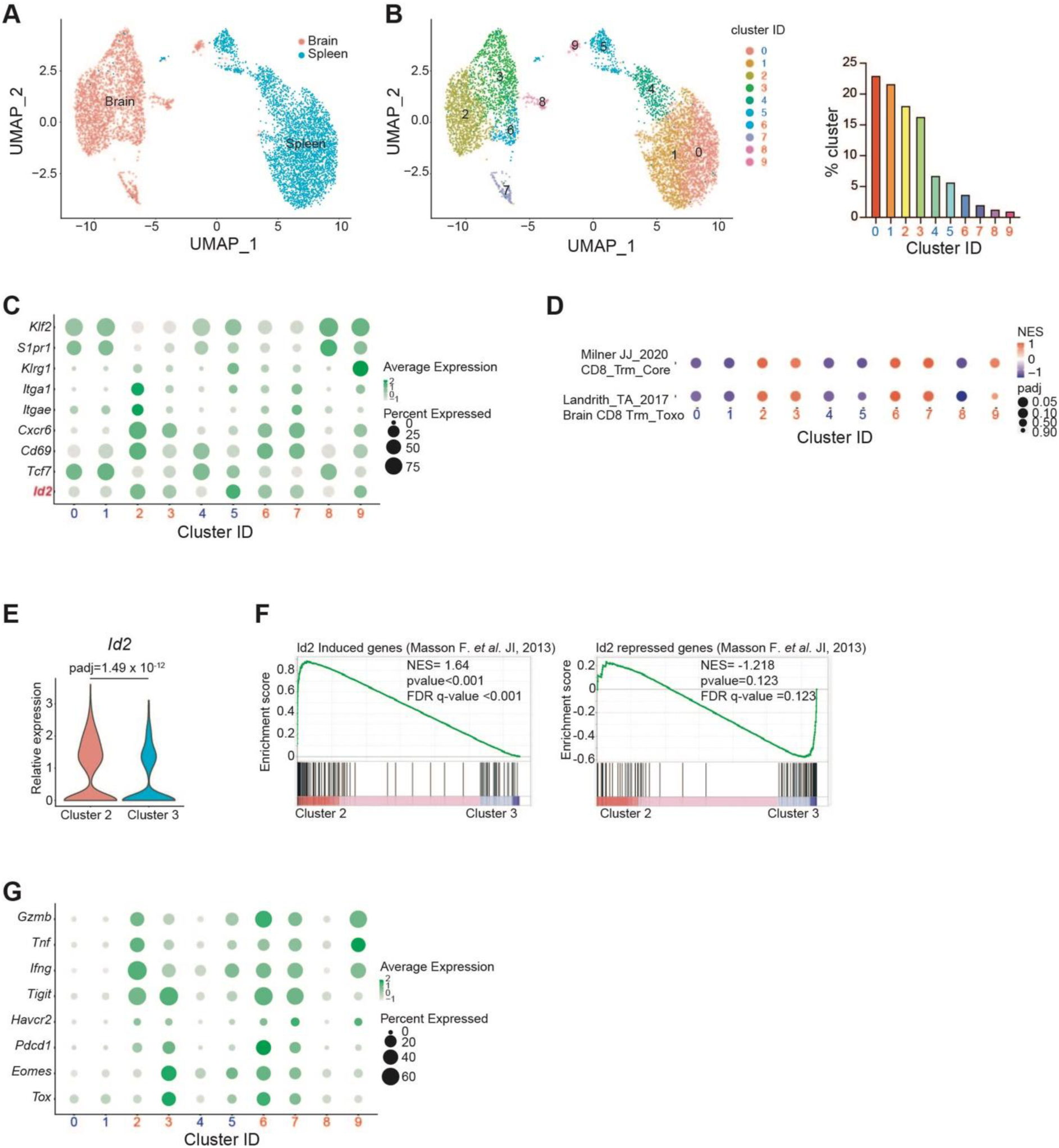
Heterogeneity of Id2 expression across brain Trm subsets correlates with their functional heterogeneity during the chronic phase of *T.gondii* infection. (A) ScRNAseq of CD8^+^ T cells isolated from the brain and the spleen of mice infected with Tg.GRA6-OVA at day 154 post-immunization. UMAP showed segregation between spleen and brain CD8^+^ T cells. (**B**) UMAP of SEURAT-generated clusters (left) and cluster frequencies (right). (**C**) Dot plot of gene expression for the indicated genes involved in CD8^+^ T cell differentiation and tissue-residency. (**D**) GSEA for the indicated published gene signatures. Normalized enrichment score (NES) and adjusted P values are shown. **(E)** Violin plot shows Id2 expression in the 2 most abundant Trm clusters Cluster 2 and Cluster 3. Statistical differences were assessed using Wilcoxon Rank Sum test. **(F)** GSEA of DE genes decreased in Id2-deficient CD8^+^ T cells (Id2-induced gene signature) or increased in Id2-deficient CD8^+^ T cells (Id2-repressed gene signature) from Masson F *et al*. 2013 (*25*) in cluster 2 vs. cluster 3. (**G**) Dot plot of gene expression for the indicated genes involved in CD8^+^ T cell exhaustion and effector functions.

Given the fact that Id2 expression has been previously shown to correlate with effector CD8^+^ T cell differentiation, we then wondered whether the heterogeneity of Id2 expression in Trm clusters could correlate with differential functionality. Accordingly, CD8^+^ T cells from cl2 displayed high expression of genes coding for effector molecules (*Ifng*, *Tnf*, *Gzmb*), high expression of *Itga1* (encoding the integrin CD49a) and low expression of exhaustion-associated genes (*Tox, Pdcd1).* Conversely, cells from cl3 exhibited a low level of effector molecules (*Ifng*, *Tnf*, *Gzmb*) combined with an “exhaustion” phenotype characterized by high expression of genes associated with exhaustion such as the transcription factors *Tox* and *Eomes* and the inhibitory receptors *Pdcd1* and *Tigit* (**Fig. 1G**). Collectively, our results indicate that Id2 expression level correlates with the functional state of brain CD8^+^ Trm during CNS chronic infection.

### Id2-deficiency leads to the formation of brain Trm with an altered phenotype during chronic CNS infection

We then endeavoured to test whether Id2 expression level actively regulates brain Trm development and function during chronic CNS *T. gondii* infection. To this end, we generated mixed bone marrow (BM) chimeras by reconstituting lethally irradiated F1 CD45.1^+^, CD45.2^+^ WT mice with a mix of BM cells from T cell-specific Id2-deficient mice Id2^fl/fl^ CD4Cre^+^ (Id2 TKO, CD45.2^+^) and BM from CD45.1^+^ WT congenic mice that were left to reconstitute 7-8 weeks before *T.gondii* infection. As additional controls for the genetic background of the Id2floxed;CD4cre colony, mixed BM chimeras reconstituted with the BM from WT Id2floxed;CD4cre littermates (noted WT CD45.2^+^ thereafter) and from WT CD45.1^+^ mice were generated. Due to an increased sensibility of the BM chimeras during the acute phase of the infection with Tg.GRA6-OVA, mice were infected with an attenuated *T.gondii* strain expressing the model antigen GRA6-OVA (Tg.ΔGRA2.GRA6-OVA) that establishes a normal latency in the brain during the chronic phase (*36*) (**Fig. 2A**). As previously reported during acute pathogen infections (*23*), Id2-deficiency in T cells strongly decreased the accumulation of pathogen-specific (K^b^OVA-dextramer(dex)^+^) CD8^+^ T cells in both brain and spleen and the loss of Id2-deficient (CD45.2^+^) OVA-specific CD8^+^ T cells was accentuated in the chronic phase (9-11 weeks post-infection) compared with the acute phase (2 weeks post-infection) of the infection (**Fig. 2B, Suppl Fig. 1A**).

**Figure 2:**
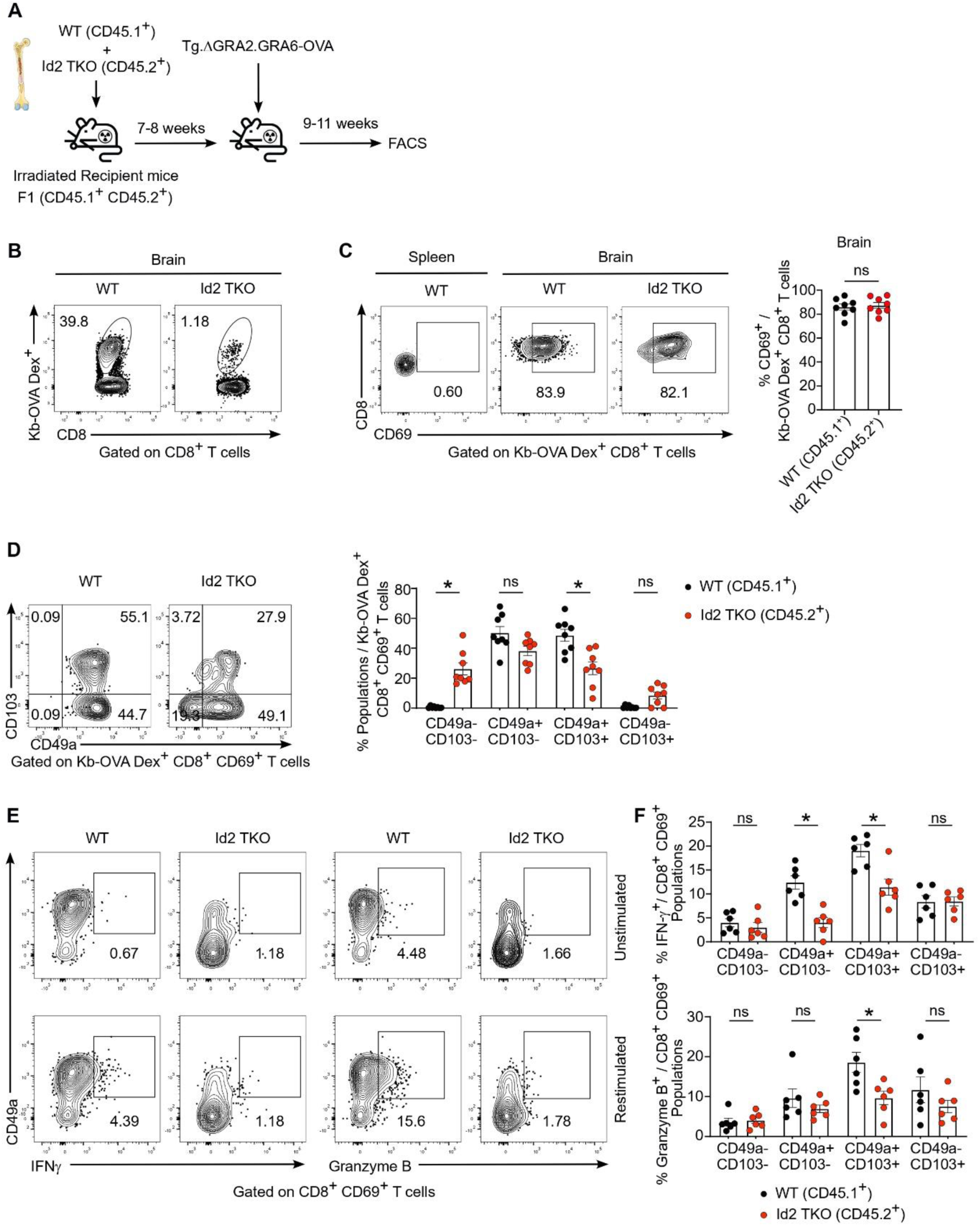
Loss of Id2 alters the differentiation of parasite-specific brain Trm during chronic *T.gondii* infection. (**A**) Experimental workflow of mixed bone marrow (BM) chimeras reconstituted with a mix of WT (CD45.1^+^) and Id2 TKO (CD45.2^+^) BM and infected with Tg.ΔGRA2.GRA6-OVA. Mice were analyzed 9-11 weeks (wks) post-infection (p.i). (**B**) Representative contour plots show the frequency of Id2 TKO (CD45.2^+^) and WT (CD45.1^+^) OVA-specific (K^b^-OVA Dex^+^) CD8^+^ T cells within the brain. (**C**) Representative contour plots show CD69 expression within K^b^-OVA Dex^+^ CD8^+^ T cells of the indicated genotype from brain and spleen. Bar graph shows the percentage of CD69^+^ cells among brain WT (CD45.1^+^) and Id2 TKO (CD45.2^+^) OVA-specific CD8^+^ T cells. (**D**) (Left) Representative contour plots of CD49a and CD103 expression among brain Id2 TKO (CD45.2^+^) or WT (CD45.1^+^) CD69^+^ K^b^-OVA Dex^+^ CD8^+^ T cells. (Right) Bar graph shows the frequency of the indicated population among brain Id2 TKO (CD45.2^+^) and WT (CD45.1^+^) CD69^+^ K^b^-OVA Dex^+^ CD8^+^ T cells. (**E**) Representative contour plots show the intracellular expression of IFN-γ and granzyme B within brain Id2 TKO and WT on CD44^+^ CD8^+^ CD69^+^ T cells after restimulation either with PMA and ionomycin in presence of monensin, or with monensin alone. (**F**) Bar graphs show the frequency of IFN-γ (Top) and Granzyme B (Bottom) expressing cells after PMA/ionomycin restimulation among total Id2 TKO or WT CD69^+^ CD8^+^ T cells. Data are pooled from 2 (D-F) to 4 (B-C) independent experiments. Bars show the mean ± SEM. Statistically significant differences were determined using Wilcoxon matched-pairs signed rank tests (C) or multiple Wilcoxon matched-pairs signed rank test corrected for multiple comparisons using Hom-Sidak method (D, F). \**P* < 0.05, \*\**P* < 0.01, \*\*\**P* < 0.001, and \*\*\*\**P* < 0.0001. ns, not significant. Each dot represents an individual mouse.

We then investigated the phenotype of OVA-specific CD8^+^ T cells during the chronic phase of *T.gondii* infection and in particular the acquisition of the Trm phenotype. Brain-infiltrating OVA-specific CD8^+^ T cells from both the Id2-deficient (CD45.2^+^) and the WT (CD45.1^+^) CD8^+^ T cell compartments of the mixed BM chimeras largely expressed the Trm marker CD69 in contrast with their splenic counterparts, and the loss of Id2 had no impact on the proportion of CD69-expressing cells compared with WT cells (**Fig. 2C**). Furthermore, WT CD69^+^ OVA-specific CD8^+^ T cells uniformly expressed CD49a and around half (48.6% ± 4.0) of CD49a^+^ cells also expressed CD103 (**Fig. 2D)**. Remarkably, compared with WT CD8^+^ T cells, Id2-deficiency led to the emergence of OVA-specific CD69^+^ CD8^+^ T cells expressing neither of these two key retention molecules, which was paralleled by a reciprocal decreased proportion of CD49a^+^ cells expressing or not CD103 (**Fig. 2D**). By contrast, control chimeras did not show any difference in Trm marker expression between the CD45.2^+^ and the CD45.1^+^ OVA-specific CD8^+^ T cell compartments **(Suppl. Fig. 1D**).

Our single-cell transcriptomic data suggested that high Id2 expression correlated with increased Trm effector functions, while lower Id2 expression was linked to a dysfunctional state. Thus, we next assessed the consequences of Id2-deficiency on effector molecules expression by brain CD8^+^ Trm cells. This analysis revealed that Id2-deficiency resulted in a marked decrease of IFN-γ expression by double positive (CD69^+^ CD49a^+^ CD103^-^) and triple positive (CD69^+^ CD49a^+^ CD103^+^) Trm, and in a significant reduction of granzyme B expression by the triple positive (CD69^+^CD49a^+^CD103^+^) Trm subset (**Fig. 2E, F**). By contrast, no difference in effector functions was found in control chimeras between the CD45.2^+^ and CD45.1^+^ WT brain CD8^+^ T cell compartments (**Suppl. Fig. 1E, F**). Altogether, our results show that Id2-deficiency leads to the emergence of parasite-specific Trm exhibiting an altered phenotype characterized by decreased expression of retention molecules and reduced effector functions (IFN-γ^+^ granzyme B^+^) during the chronic phase of *T. gondii* infection.

### Id2-mediated inhibition of E2A is required for CD49a^+^ effector Trm differentiation

We reasoned that the decreased frequency of CD49a^+^ effector Trm upon Id2-deficiency could either reflect a loss of this population over time, or instead could suggest that Id2 played a role in the differentiation of this Trm subset. To resolve this issue, we analysed the early development of Trm during the acute phase of the infection, i.e. 2 weeks after Tg.ΔGRA2.GRA6-OVA inoculation. At this early time-point, a similar proportion of WT and Id2-deficient OVA-specific CD8^+^ T cells expressed CD69 (**Fig. 3A)**. Importantly, akin to the chronic phase, we observed a decreased frequency of CD49a-expressing cells and the reciprocal increase in double negative cells among Id2-deficient OVA-specific CD69^+^ CD8^+^ T cells (**Fig. 3B, C**). As expected, CD103 was only weakly expressed at this early stage of the infection (**Fig. 3C**). Moreover, granzyme B expression was downregulated in total CD8^+^ CD69^+^ Trm cells during the acute phase of the infection, albeit to lower extent compared with the chronic phase **(Fig. 3D**). Taken together, these results suggested that Id2 loss cripples the differentiation of effector Trm subtypes rather than impedes their maintenance.

**Figure 3:**
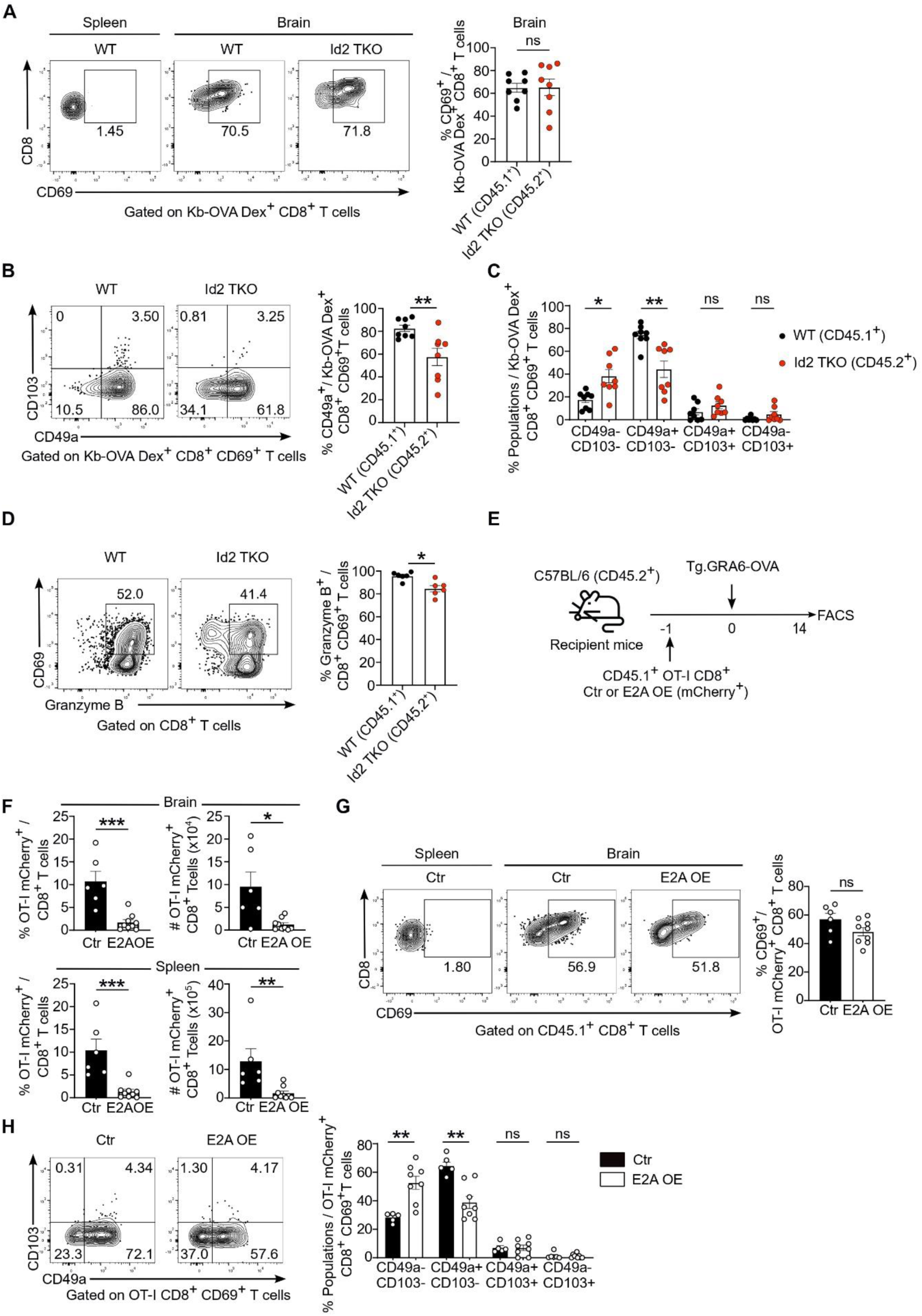
The transcriptional axis Id2-E2A regulates the early differentiation of CD49a^+^ brain Trm during the acute phase of *T.gondii* infection. (A-D) Mixed BM chimeras Id2 TKO (CD45.2^+^) :WT (CD45.1^+^) were generated as described in Fig.2A and were analyzed during the acute phase (2 wks p.i.) of Tg.ΔGRA2.GRA6-OVA infection. (**A**) Representative contour plot (left) and bar graph (right) show the frequency of CD69^+^ cells among brain Id2 TKO (CD45.2^+^) and WT (CD45.1^+^) K^b^-OVA Dex^+^ CD8^+^ T cells. (**B**) Representative contour plot (left) and bar graph (right) show the frequency of CD49a^+^ cells among brain Id2 TKO (CD45.2^+^) and WT (CD45.1^+^) K^b^-OVA Dex^+^ CD69^+^ CD8^+^ T cells. (**C**) Bar graph shows the percentage of the indicated CD8^+^ T cell population among brain Id2 TKO (CD45.2^+^) and WT (CD45.1^+^) K^b^-OVA Dex^+^ CD69^+^ CD8^+^ T cells. (**D**) (Left) Representative contour plots show granzyme B expression among brain Id2 TKO (CD45.2^+^) and WT (CD45.1^+^) total CD44^+^ CD8^+^ T cells and (right) bar graph shows the frequency of granzyme B expressing cells among brain Id2 TKO and WT CD69^+^ CD44^+^ CD8^+^ T cells. (**E**) Schematic diagram outlining the adoptive transfer of TCR transgenic OVA-specific CD8^+^ T cells (OT-I) transduced with retroviruses encoding the E2A isoform E47 (E2A OE) or a control (Ctr) empty vector and the mCherry as fluorescent reporter, to recipient mice subsequently infected with Tg.GRA6-OVA. Mice were analyzed at day 14 p.i and brains were collected. (**F**) Bar graphs show the frequency among CD8^+^ T cells (left) and total number (right) of Ctr and E2A OE OT-I mCherry^+^ (CD45.1^+^) in the brain and spleen. (**G**) Representative contour plots (left) and bar graph (right) show the percentage of brain CD69^+^ cells among Ctr and E2A OE OT-I mCherry^+^ (CD45.1^+^) cells. (**H**) (Left) Representative contour plots show the expression of CD103 vs. CD49a in Ctr and E2A OE in brain CD69^+^ OT-I mCherry^+^ (CD45.1^+^) CD8^+^ T cells. (Right) Bar graph shows the percentage of the indicated population within Ctr and E2A OE CD69^+^ OT-I mCherry^+^ (CD45.1^+^) cells from the brain. Data are pooled from 2 (D, F, G, H) and 3 (A, B, C) independent experiments. Bars show the mean ± SEM. Statistically significant differences were determined using multiple Wilcoxon matched-pairs signed rank tests corrected for multiple comparisons using Hom-Sidak method (C), Paired t-test (A, B, D), Mann-Whitney (F, G) and multiple Mann-Whitney (H) tests corrected for multiple comparisons using Hom-Sidak method. \**P* < 0.05, \*\**P* < 0.01, \*\*\**P* < 0.001, and \*\*\*\**P* < 0.0001. ns, not significant. Each dot represents an individual mouse

Id2 has been previously shown to repress memory precursor formation through the inhibition of E2A transcriptional activity during acute pathogen infection (*25*). To determine whether Id2 could also act through the inhibition of E2A activity during Trm development, we asked whether E2A overexpression (OE) could phenocopy the impact of Id2-deficiency on the Trm phenotype. To address this point, OVA-specific OT-I CD8^+^ T cells were transduced with a retroviral vector encoding E2A (E47 isoform) and mCherry as reporter, or with a control-IRES-mCherry vector and were transferred into recipient mice that were infected with Tg.GRA6-OVA one day later (**Fig 3E**). Two weeks post-infection, the total number and frequency of OT-I cells overexpressing E2A were drastically decreased compared with control OT-I cells both in brain and in spleen (**Fig 3F**). Phenotypically, E2A OE did not impact the proportion of CD69 expressing OT-I cells infiltrating the brain but it led to a significant decreased proportion of CD49a^+^ OT-I cells among CD69^+^ cells compared with OT-I transduced with the control vector, similarly to the effect observed upon Id2-deficiency (**Fig. 3G-H**). These results suggested that Id2 promoted the formation of the CD49a^+^ Trm subsets through the inhibition of E2A.

Since Id2 has been reported to act in a dose-dependent manner during peripheral CD8^+^ T cell differentiation to an acute viral infection (*25*), we next investigated whether increasing Id2 dosage could enhance the accumulation of OT-I cells and the generation of brain Trm cells expressing CD49a during Tg.GRA6-OVA infection (**Suppl. Fig. 2A**). Remarkably Id2 OE strongly increased the total number of OT-I cells in both brain and spleen (**Suppl. Fig. 2B-C**). Moreover, in contrast to E2A OE, the frequency of CD49a^+^ expressing cells among CD69^+^ OT-I Trm increased upon Id2 OE (**Suppl. Fig. 2D, E).**

Next, we explored the molecular mechanisms underpinning this increased formation of CD49a^+^ Trm cells upon Id2 OE. As the expression of the transcriptional regulator Hobit has been reported to correlate with the expression of CD49a in Trm (*37*), we hypothesized that Id2 may promote the generation of CD49a^+^ Trm through the induction of Hobit expression. To address this question, we made use of OT-I cells crossed with the Hobit-TdTomato-Cre-DTR reporter mouse (*38*) to test the effect of Id2 OE on Hobit expression in developing brain OT-I Trm during the acute phase of *T. gondii-*OVA infection. Our data showed that the expression level of Hobit-TdTomato in various CD69^+^ Trm subsets (CD49a^-^CD103^-^, CD49a^+^CD103^-^, CD49a^+^CD103^+^) was not impacted by Id2 OE, indicating that Id2 OE do not increase the formation of CD49a-expressing Trm through the increase of Hobit expression (**Suppl. Fig. 2F,-G**).

### Id2-deficiency promotes the generation of brain Trm exhaustion during chronic *T. gondii* infection

Our transcriptomic data of CD8^+^ T cells isolated from the brain of chronically *T.gondii* infected mice showed that the cluster of Trm exhibiting a lower Id2 expression (cl3) displayed a profile of T cell exhaustion characterized by high *Tox* and *Pdcd1* expression (see **Fig. 1G**). These observations led us to hypothesize that the downregulation of Id2 expression might promote Trm exhaustion during chronic pathogen infection. To address this issue, we analysed the exhaustion phenotype of CD8^+^ T cells isolated from the brain of Id2 TKO (CD45.2^+^):WT (CD45.1^+^) mixed BM chimeras and WT (CD45.2^+^):WT (CD45.1^+^) control chimeras during the acute and chronic phases of the infection. In line with former observations in the *T.gondii* infection model (*5, 39*) both brain-infiltrating WT and Id2-deficient parasite-specific (K^b^OVA Dex^+^)-specific CD69^+^ CD8^+^ T cells did not exhibit any sign of exhaustion during the chronic stage of the infection, as shown by their lack of PD-1 and Tox expression (**Fig. 4A, B**). Instead, cells co-expressing PD-1 and Tox were found among K^b^OVA-Dex negative brain-infiltrating CD69^+^ CD8^+^ T cells (**Fig. 4A, B**). This population of CD8^+^ T cells developed specifically during the chronic stage of the infection as indicated by the absence of PD-1 and Tox expression during the acute phase of the infection (**Fig. 4B**). Remarkably, the proportion of cells expressing Tox and PD-1 among K^b^OVA-Dex negative CD8^+^ T cells significantly increased among Id2-deficient CD8^+^ T cells compared with WT CD45.1^+^ CD8^+^ T cells from Id2 TKO:WT mixed chimeras, while no differences were found between WT CD45.2^+^ and WT CD45.1^+^ CD8^+^ T cells in control chimeras (**Fig. 4B** & **Suppl. Fig. 3A, B**). This increased proportion of PD-1^+^ Tox^+^ cells was found across all Trm subsets, although the magnitude of the effect was higher in CD69^+^ CD49a^+^ CD103^-^ cells (**Fig. 4C**).

**Figure 4:**
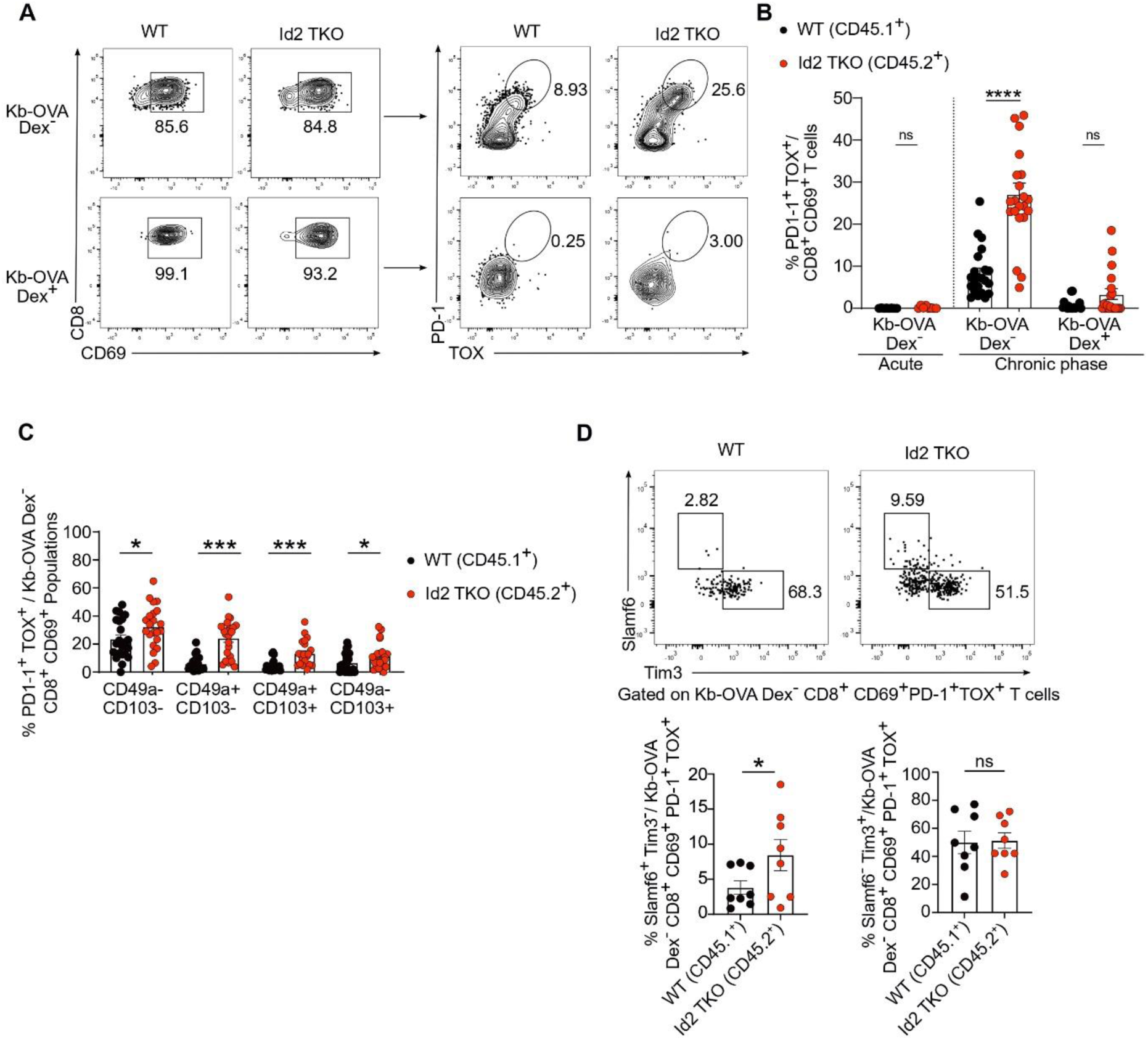
Id2-deficiency leads to increased proportion of exhausted CD8^+^ T cells in the brain of mice chronically infected with *T.gondii*. Mixed BM chimeras Id2 TKO (CD45.2^+^) :WT (CD45.1^+^) were generated as described in Fig. 2A and were analyzed during the acute phase (2 wks p.i.) and the chronic phase (9-11 wks p.i.) of Tg.ΔGRA2.GRA6-OVA infection. **(A)** Gating strategy and representative contour plots of PD-1 and Tox expression in WT (CD45.1^+^) and Id2 TKO (CD45.2^+^) K^b^-OVA Dex negative CD8^+^ CD44^+^ CD69^+^ T cells (top) and K^b^-OVA Dex^+^ CD8^+^ CD69^+^ T cells (bottom) from the brain during the chronic phase of the infection. (**B**) Bar graph shows the frequency of PD-1^+^TOX^+^ expressing cells either among WT (CD45.1^+^) and Id2 TKO (CD45.2^+^) K^b^-OVA Dex^+^ CD69^+^ CD8^+^ T cells or among K^b^-OVA Dex negative CD8^+^ CD44^+^ CD69^+^ T cells from the brain during the acute and chronic stage of the infection. (**C**) Bar graph shows the percentage of PD-1^+^TOX^+^ expressing cells within the indicated population among brain WT (CD45.1^+^) and Id2 TKO (CD45.2^+^) K^b^-OVA Dex negative CD69^+^ CD44^+^ CD8^+^ T cells during the chronic phase of the infection. (**D**) Representative dot plots and bar graph show the proportion of Slamf6^+^Tim3^-^ and PD1-Slamf6^-^Tim3^+^ cells among Id2 TKO (CD45.2^+^) and WT (CD45.1^+^) PD-1^+^ Tox^+^ CD69^+^ CD44^+^ CD8^+^ T cells. Data are pooled from 2 (D) and 6 (B, C) independent experiments. Bars show the mean ± SEM. Statically significant differences were determined using Paired t-test (D), Wilcoxon matched-pairs signed rank test (B) and multiple Wilcoxon matched-pairs signed rank test corrected for multiple comparisons using Hom-Sidak method (C). \**P* < 0.05, \*\**P* < 0.01, \*\*\**P* < 0.001, and \*\*\*\**P* < 0.0001. ns, not significant. Each dot represents an individual mouse.

Tox has been previously reported to repress Id2 expression and Id2-dependent gene signature in encephalitogenic T cells in a model of autoimmune encephalitis (*33*). Since Id2 is important to repress E2A-mediated activation of *Tcf7 (*encoding Tcf1) expression during an acute viral infection (*25*), and that Tcf1 is required for the generation of Tpex during chronic infection, we then investigated whether Id2-deficiency may impact the proportion of Tox^+^ CD8^+^ T cells exhibiting Tpex features. To address this issue, we investigated the expression of Slamf6 (a surrogate of Tcf1 expression) and Tim-3 in brain infiltrating PD1^+^ Tox^+^ CD69^+^ CD8^+^ T cells. Remarkably, Id2-deficiency led to a significant increase of the relative proportion of the Slamf6^+^Tim-3^-^ Tpex-like subset, while the frequency of Slamf6^-^Tim3^+^ terminally exhausted cells was not impacted (**Fig. 3D**). Finally, no difference was noted between the CD45.1^+^ and CD45.2^+^ WT Tox^+^ PD-1^+^ CD69^+^ CD8^+^ T cell compartments from control chimeras (**Suppl. Fig. 3C)**. Taken together, our data indicated that Id2-deficiency increased the proportion of exhausted CD8^+^ Trm cells and altered their balance of differentiation between a Tpex-like phenotype and a terminal exhaustion phenotype during the chronic phase of *T. gondii-*OVA infection.

### Id2-forced expression represses Trm exhaustion during chronic *T. gondii* infection

Given that our data had shown an increased frequency of exhausted CD8^+^ Trm cells upon loss of Id2, we then wondered whether increasing Id2 level could repress brain Trm cell exhaustion and rewire the program of exhaustion towards an effector program. Since our model of *T. gondii*-OVA chronic infection yielded only a modest level of CD8^+^ T cell exhaustion and that parasite-specific CD8^+^ T cells were protected from exhaustion (**Fig. 4A**), we wished to set out a second model of *T. gondii* infection leading to an increased level of exhaustion in brain CD8^+^ T cells, including among parasite-specific CD8^+^ T cells. Because CD4^+^ T cell help has been previously shown to repress CD8^+^ T cell exhaustion during *T. gondii* infection (*40*), we tested the effect of CD4^+^ T cell depletion on parasite-(OVA)-specific CD8^+^ Trm cell exhaustion. As anticipated, CD4^+^ T cell depletion led to an increased level of T cell exhaustion in K^b^OVA-Dex negative CD44^+^ CD8^+^ T cells but also among brain-infiltrating K^b^OVA-Dex^+^ parasite-specific CD8^+^ T cells 35 days post-infection (**Fig. 5A**). Using this experimental setting, we next investigated the impact of Id2 OE on the accumulation and the exhaustion of brain-infiltrating OVA-specific OT-I CD8^+^ T cells (**Suppl. Fig. 4A**). Strikingly, OT-I cells overexpressing Id2 strongly outnumbered OT-I cells transduced with the control vector both in spleen and brain 5 weeks post-infection (**Fig. 5B, C**). Brain infiltrating OT-I cells from both groups largely adopted a Trm phenotype co-expressing CD69 and CD49a at this chronic stage of the infection (**Suppl. Fig. 4B, C**). However, Id2 OE reduced the proportion of cells co-expressing PD-1 and Tox among CD69^+^ OT-I cells (**Fig. 5D**). The parasite load in the brain was similar between mice transferred with control and Id2 OE OT-I cells, as well as the proportion of exhausted cells among the endogenous brain CD8^+^ T cell compartment (**Suppl. Fig. 4D, E**). This indicated that the reduced exhaustion of Id2 OE OT-I cells was intrinsically caused by the increased level of Id2 expression and was not due to differences in antigen load or inflammation, which would have seemingly impacted the phenotype of the endogenous CD8^+^ T cell compartment. Analysis of the proportion of terminally exhausted vs Tpex-like cells among total PD-1^+^ Tox^+^ CD69^+^ exhausted OT-I cells revealed a reduction of the frequency of the Tpex-like subset (Slamf6^+^ Tim3^-^) upon Id2 OE (**Fig. 5E**) in line with our previous data showing that instead, Id2-deficiency increased the frequency of this subset (**Fig. 4C**). T cell exhaustion is characterized by a diminished capacity to co-produce multiple effector cytokines such as IFN-γ and TNF. Our results showed that brain Id2 OE OT-I Trm (CD69^+^) had an increased proportion of total IFN-γ producers and IFN-γ/TNF double producers consistent with the induction of an effector-like phenotype and a decreased functional exhaustion (**Fig. 5F)**.

**Figure 5:**
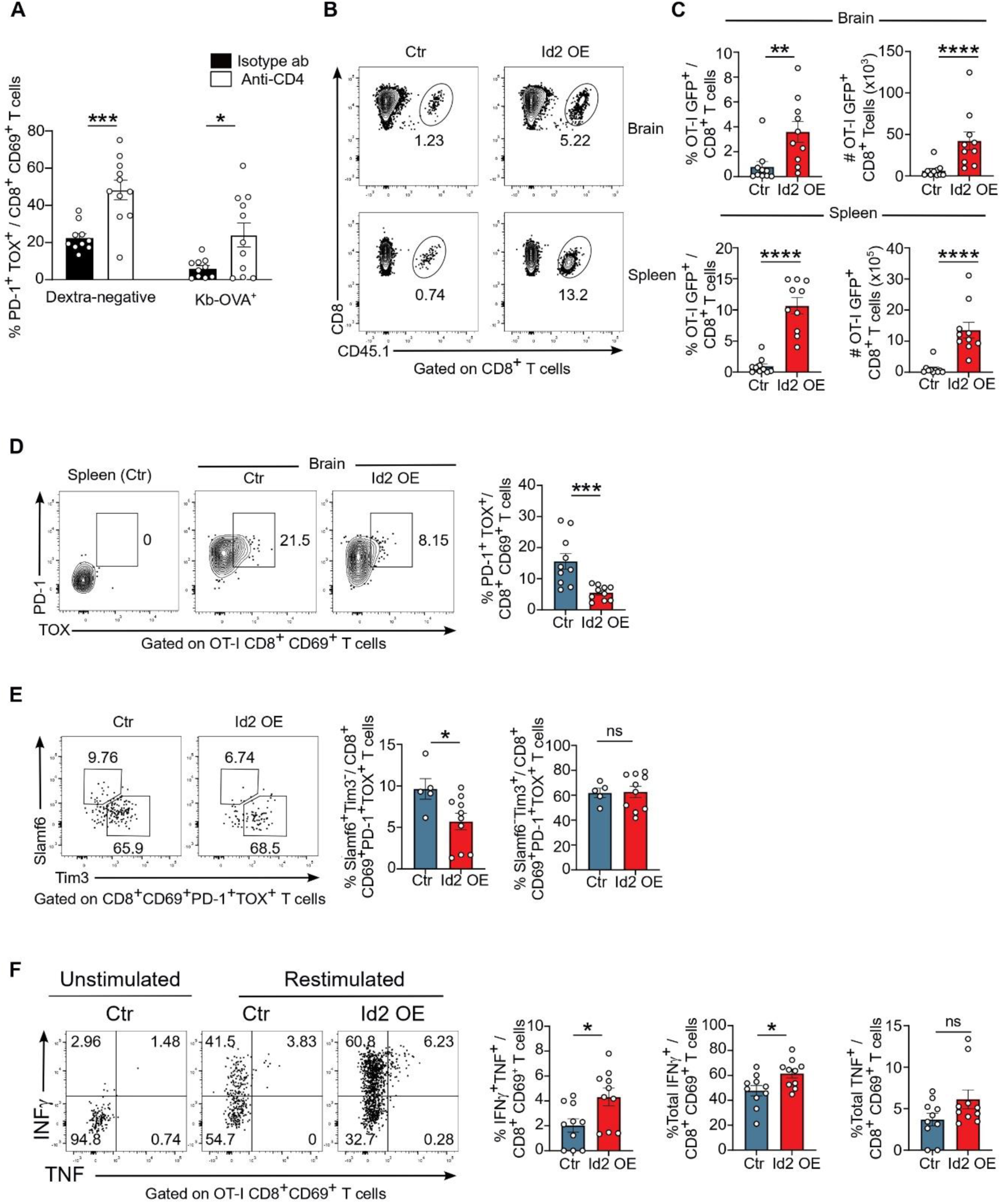
Id2 overexpression represses the exhaustion of parasite-specific Trm during the chronic phase of *T. gondii* upon CD4^+^ T cell depletion. (**A)** Mice received weekly injection either an anti-CD4 depleting antibody or an isotype control antibody from day-3 to day 35 p.i. with Tg.GRA6-OVA. Mice were analyzed 5 wks p.i. and brain and spleen were collected. Bar graph shows the proportion of PD-1^+^TOX^+^ expressing cells in K^b^-OVA Dex^-^ or Dex^+^ CD8^+^ T cells from the brain at the chronic stage. (**B-F**) Mice were adoptively transferred with OT-I (CD45.1^+^) cells transduced with a control vector or an Id2 overexpressing (Id2 OE) vector expressing the GFP reporter one day prior Tg.GRA6-OVA infection. All mice were treated with the anti-CD4 depleting antibody as in **A.** (**B**) Representative contour plots show the percentage of Ctr and Id2 OE OT-I GFP^+^ among total CD8^+^ T cells from the brain (top) and spleen (bottom). (**C**) Bar graphs show the frequency among CD8^+^ T cells (left) and total number (right) of Ctr and Id2 OE OT-I GFP^+^ (CD45.1^+^) in the brain and spleen. (**D**) Representative contour plots and bar graphs show the percentage of PD-1^+^TOX^+^ cells among brain CD69^+^ Id2 OE and Ctr OT-I (CD45.1^+^) cells. (**E**) Dot plots and bar graphs show the frequency of Slamf6^+^Tim3^-^ and Slamf6^-^Tim3^+^ cells among Id2 OE and Ctr PD-1^+^ Tox^+^ CD69^+^ OT-I (CD45.1^+^). (**F**) Dot plots and bar graphs show the frequency of IFN-γ^+^ and TNF^+^ producing cells in brain Id2 OE and Ctr CD69^+^ OT-I (CD45.1^+^) cells after restimulation with OVA peptide in presence of monensin, or with monensin alone as control. Data are pooled from 2 independent experiments. Bars show the mean ± SEM. Statistically significant differences were determined using unpaired t-test (A) and Mann-whitney test (C-F). \**P* < 0.05, \*\**P* < 0.01, \*\*\**P* < 0.001, and \*\*\*\**P* < 0.0001. ns, not significant. Each dot represents an individual mouse.

The decreased level of T cell exhaustion observed upon Id2 OE during *T.gondii* chronic infection could be explained either by a direct impact of Id2 OE to repress the program of T cell exhaustion or instead, it could reflect an Id2-dependent preferential accumulation of Trm with an effector phenotype over the 5 weeks of infection. Thus, we next asked whether Id2 OE intrinsically represses the acquisition of the exhaustion phenotype. To this end, we adapted a previously published protocol of *in vitro* CD8^+^ T cell exhaustion based on repeated antigenic stimulation (*41*) (**Suppl. Fig. 5A)**. After 7 days of OVA peptide stimulation, most control and Id2 OE OT-I upregulated PD-1 and a large proportion of them co-expressed Tox, while cells stimulated only once with OVA peptide failed to upregulate these exhaustion markers (**Suppl. Fig. 5B**). Strikingly, Id2 OE led to a decreased frequency of PD-1^+^Tox^+^ co-expressing cells compared with the control transduced cells akin to our observations made in the *T. gondii* chronic infection model *in vivo* (**Suppl. Fig. 5B-D**). In addition, Id2 OE partially restaured effector functions with an increased proportion of granzyme B and IFN-γ expressing cells, and improved polyfunctionality as shown by the increased proportion of IFN-γ/TNF co-producing cells (**Suppl. Fig. 5E-G**). Collectively, our data showed that forced-Id2 expression inhibits T cell exhaustion during chronic *T. gondii* infection and after *in vitro* repeated antigenic stimulation, highlighting its potential as a therapeutic target for maintaining T cell functionality in persistent infections and cancer.

## DISCUSSION

Recent single-cell transcriptomic studies have revealed that Trm are a very heterogenous T cell population during infection, autoimmunity and tumor development in both humans and mice (*5, 10–14*). However, the molecular determinants regulating this phenotypic and functional diversity are not fully understood. In a model of chronic CNS infection with the intracellular parasite *T. gondii*, we demonstrated that Id2 expression levels and its interaction with E2A in pathogen-specific CD8^+^ T cells regulate the balance between the differentiation of effector Trm, which express high levels of effector molecules and of the tissue retention molecule CD49a, and dysfunctional brain Trm cells, which exhibit an exhausted phenotype and reduced effector functions. Using single-cell transcriptomic of brain and splenic CD8^+^ T cells isolated from *T.gondii* chronically infected mice, we found that *Id2* expression was more elevated in the Trm cluster showing the highest level of effector molecules expression. By contrast, among the two most abundant brain Trm clusters identified in our study, the cluster of cells showing the lowest *Id2* expression exhibited an exhausted phenotype, characterized by high expression of *Tox*, *Pdcd1*, and *Tigit,* and low levels of effector and cytotoxic molecules. Using mixed BM chimeras, we showed that T-cell specific Id2-deficiency led to a decreased development of parasite-specific brain Trm cells expressing CD49a and CD103, and to reduced effector cytokines and cytotoxic molecule expression. Moreover, the Id2-deficient CD8^+^ T cell compartment was enriched in Trm cells displaying an exhausted phenotype characterized by Tox and PD-1 expression.

Interestingly, a previous meta-analysis of single-cell RNAseq data of CD8^+^ T cells from the CSF and brain of multiple sclerosis (MS) patients has defined a common CD4^+^ and CD8^+^ Trm minimal gene signature that included ID2, suggesting that ID2 could also be important for the generation of CNS-resident Trm in humans (*13*). It remains unclear whether Id2 plays a specific role in the differentiation of brain Trm or if it more broadly contributes to the differentiation of Trm in other anatomical sites. However, the initial description of the Id2 KO mice and recent data in acute LCMV infection have shown that Id2-deficiency led to a loss of IEL of the intestine hinting towards a global role of Id2 in Trm development, although the mechanisms underpinning this loss have not been fully explored (*11, 27*). Here, our study revealed that, besides its reported role in intestinal Trm homeostasis, Id2 impacts the phenotype and the functions of brain Trm. Indeed, loss of Id2 led to reduced frequencies of CD69^+^ T cell subsets expressing the tissue-retention molecule CD49a alone or co-expressed with CD103, and instead caused the emergence of atypical Trm cells expressing neither of these integrins. Importantly E2A OE in pathogen-specific CD8^+^ T cells resulted in a similar deregulation of the expression of CD49a during early Trm development. This suggested that Id2 antagonizes E2A transcriptional activity to enable the formation of fully functional Trm capable of long-term retention within the brain. Indeed, CD49a has been shown to be required for the long-term retention of Trm in non lymphoid tissues such as the skin (*42*). Moreover, decreased CD49a expression could also potentially contribute to the decreased effector functions observed upon Id2-deficiency. Indeed, previous studies have reported that CD49a-deficient Trm cells had reduced IFN-γ expression compared with CD49a-sufficient cells (*42*). Similarly, CD49a^+^ CD103^+^ Trm from human psoriatic skin lesions have been shown to possess an increased cytotoxic potential compared with their CD49a-negative counterparts (*43*).

Id2, together with Blimp-1 and T-bet, are key transcriptional regulators of effector CD8^+^ T cell differentiation (*19–21, 23–26*). Interestingly, Blimp-1 and T-bet not only promote the acquisition of an effector phenotype but also have been shown to regulate Trm differentiation and T cell exhaustion during infection (*17, 18, 44, 45*). Indeed, while Blimp-1 and its homologue Hobit are required for Trm differentiation (*17*), the downregulation of T-bet (along with Eomes) is necessary to permit TGF-β signaling crucial for Trm development (*18*). Here, our data showing a role for Id2 in Trm functions reinforce the concept that the effector and the tissue-residency transcriptional programs of CD8^+^ T cell differentiation are tightly interconnected and dependent on a similar set of transcriptional regulators. Moreover, Blimp-1 and T-bet have also divergent roles in the development of terminally exhausted cells since T-bet deficiency exacerbates T cell dysfunction and increases inhibitory receptor expression, (*45*), whereas Blimp-1 deficiency abrogates terminal T cell exhaustion in a dose-dependent manner (*44*). Interestingly, both Blimp-1 and Id2 T-cell deficiencies have been previously shown to enhance the differentiation of splenic “follicular cytotoxic CD8^+^ T cells” during chronic LCMV infection. These cells exhibited a phenotype reminiscent of Tpex, characterized by the co-expression of PD-1 and TCF-1 (*30*). This is consistent with our results showing that Id2 loss led to an increase proportion of Tox^+^ PD-1^+^ exhausted Trm cells displaying a Tpex-like phenotype.

Conversely, Id2 OE promoted CD8^+^ T cell accumulation, decreased Tox and PD-1 expression and led to some recovery of effector functions both during chronic *T.gondii* infection and in an *in vitro* model of T cell exhaustion. These results suggested that the exhaustion program of CD8^+^ T cell differentiation can be rewired by enforcing the effector program through the forced expression of Id2. This finding is in line with recent studies, which have reported that activated STAT5 OE or IL-2 therapy could promote an “effector-like” phenotype in exhausted T cells and repress the Tox-dependent epigenetic and transcriptional program of exhaustion (*46–48*). Interestingly, pSTAT5 is a direct transcriptional activator of Id2 (*26*), and Id2 and STAT5 have been shown to be downregulated in a Tox-dependent manner during CNS autoimmunity and chronic viral infection, respectively (*33, 46*). This raises the question of whether the decrease in T cell exhaustion reported upon IL-2 therapeutic administration or after STAT5 forced expression (*46–48*) could be in part due to the induction of Id2 expression.

In line with previous studies (*5, 39*), our data confirmed the absence of exhaustion of parasite-specific CD8^+^ T cells infiltrating the brain during chronic *T.gondii* infection. By contrast, systemic CD4^+^ T cell depletion led to a marked increase in CD8^+^ T cell exhaustion among OVA-specific CD8^+^ T cells consistent with previous reports showing the importance of CD4 T cell help in repressing T cell exhaustion during various chronic pathogen infections (*49–52*). As CD4^+^ T cells have the potential to produce cytokines that are able to induce Id2 in CD8^+^ T cells such as IL-2 or IL-21 (*26, 53*), it is tempting to speculate that parasite-specific CD8^+^ T cells receive a sufficient amount of CD4^+^ T cell help to optimally induce Id2, which in turn represses T cell exhaustion. By contrast, the PD1^+^ Tox^+^ CD8^+^ T cell population identified among the non-OVA-specific CD8^+^ T cell population may represent CD4 “helpless” CD8^+^ T cells. It would be worthy to determine whether these PD1^+^ Tox^+^ T cells are clonally expanded and responding to parasite-derived antigens, or instead represent a bystander polyclonal CD8^+^ T cell subset.

In summary, our study has identified Id2 as a central regulator of Trm differentiation and function during chronic brain pathogen infection and suggested that Id2 expression level determines the accumulation and the functional fate of brain Trm into either “effector” Trm or “exhausted” Trm. Targeting Id2-E protein transcriptional axis could therefore be beneficial to improve Trm survival and effector functions not only in case of chronic pathogen infection but also during tumor development.

## MATERIAL AND METHODS

### Mice

C57BL/6J CD45.2^+^ (Janvier-Labs), Id2 floxed (*25*) x CD4cre mice, C57BL/6 CD45.1 (*Ptprc*) x CD45.2 mice, OT-I TCR transgenic x C57BL/6 CD45.1 (*Ptprc*) and OT-I x Hobit-TdTomato-Cre-DTR mice (*38*) were bred in house at the US006 animal facility. Recipient mice used for OT-I cell adoptive transfer experiments were all males 7-8 weeks old. All mice were bred and maintained under specific pathogen-free conditions in accordance with the guidelines of the French National Veterinary Services and the European regulations (EEC Council Directive, 2010/63/EU, September 2010) after the approval of the local ethics committee.

### Toxoplasma gondii culture and infection

For all experiments, GFP^+^ type II Prugnaud (Pru) *T. gondii*, expressing GRA6-OVA under the control of the endogenous GRA6 promoter (Pru.GFP.GRA6-OVA (*34*) or its derivative Pru.ΔGRA2.GRA6-OVA (*36*) were used. The GRA2-deficient Pru.ΔGRA2-GRA6-OVA parasites were engineered from the parental Pru.GFP.GRA6-OVA by CRISPR/Cas9-mediated suppression of the GRA2 gene, using phleomycin selection. In brief, Pru.GFP.GRA6-OVA tachyzoites were co-transfected by electroporation (1:1 ratio) with one plasmid coding for both Cas9 and a sgRNA targeting GRA2 (TGGT1_227620 gene) constructed from pSAG1::CAS9-U6::sgUPRT (*54*), by replacing the UPRT-targeting sgRNA by a sgRNA targeting GRA2: sequence 5’-TTTTCCGGAGTTGTTAACCA, corresponding to sgRNA TGGT1_227620_5 described in Sidik *et al.* (*55*) and the pGRA2/Ble/GRA2 plasmid coding for the bleomycin hydrolase resistance marker flanked by 5′ and 3’ UTR of GRA2 gene (*56*). Selection was done by a 3h pulse with 50 μg/ mL phleomycin followed by subsequent culture with 5 μg/ mL phleomycin final concentration. All tachyzoites were cultured *in vitro* through successive passages on confluent monolayers of human foreskin fibroblasts (HFF) cells using DMEM enriched with 1% FCS (GIBCO). HFF were acquired from ATCC (Hs27 ref # CRL-1634). For mouse infections, infected HFF were scraped, tachyzoites were released through a 23G needle, filtered through a 3 µm polycarbonate hydrophilic filter (it4ip S.A.) and 200 Tg.GRA6-OVA or 1000 Tg.ΔGRA2-GRA6-OVA tachyzoites were injected intra-peritoneally (i.p.) in 200 µl PBS. In mice adoptively transferred with OT-I T cells, *T. gondii* infection was performed one day after the injection of OT-I T cells.

### Generation of bone marrow chimeras

Mixed bone marrow chimeras Id2 TKO (CD45.2^+^): WT (CD45.1^+^) were generated by transplanting a mixture of wild-type (CD45.1^+^) and Id2^fl/fl^CD4^Cre+^ (CD45.2^+^) bone marrow (BM) cells at a ratio 1 : 4 and control mixed chimeras were produced using a mixture of BM cells from WT (CD45.1^+^) mice and bone marrow cells from Id2^fl/+^CD4^Cre-^ or Id2^+/+^ CD4Cre^+^ (noted as WT CD45.2^+^) littermates from the Id2floxed;CD4cre colony at a ratio 1 : 2 to lethally irradiated (800 rads) CD45.1^+^ CD45.2^+^ (F1) recipient mice. Mice were allowed to reconstitute for 7-8 weeks before infection.

### Isolation and adoptive transfer of OT-I CD8^+^ T cells

TCR transgenic OVA-specific CD8^+^ T cells (OT-I) were isolated from spleen and inguinal lymph nodes (LNs) of TCR transgenic mice by negative selection using the Dynabeads Untouched Mouse CD8 Cells kit (Invitrogen). Naive CD8^+^CD44^-^ T lymphocytes were subsequently isolated using the CD8α T Cell Isolation Kit (Miltenyi biotec) through addition of an anti-CD44 biotinylated antibody (clone IM7, BD pharmingen) to the antibody mix by negative selection using an MS column and MidiMACS separator. CD8^+^ T cells were then cultured at 1.10^6^ cells/well in 12 well/ plate that were coated overnight one day earlier with anti-CD3 mAb (clone145-2C11, BD pharmingen, 5 µg/ml) within 2 ml/well of complete medium (RPMI 1640, 10% FBS (Gibco), 1% 2mM L-glutamine (Life Technologies), 1% HEPES (Life Technologies), 1% non-essential amino acids (Life Technologies), 100U/ml penicillin (Gibco) and 100μg/ml Streptomycin-sulfate (Gibco), 0.05mM βmercaptoethanol (Sigma) supplemented with anti-CD28 mAb (clone 37.51, BD pharmingen, 2 µg/ml) and recombinant human IL-2, mouse IL-7 and IL-15 (5ng/ml) (all from Peprotech) for 3 days. Cells were transduced with retroviruses 24h after activation and 10.10^3^ cells of FACS-sorted retrotransduced cells were intravenously transferred into recipient mice 3 days later.

### Retroviral transduction of OT-I TCR transgenic T cells

Murine stem cell virus (MSCV)-Id2-IRES-GFP, MSCV-IRES-GFP, MSCV-Tcfe2a-IRES-Cherry and MSCV-IRES-Cherry have been previously described (*25, 57*). OT-I CD8^+^ T cells were enriched from the spleen and inguinal LNs as described above. Retroviral supernatants were then spun for 1 h at 3100g at 4 °C in 12-well-plates coated with 32 mg/ml RetroNectin (Takara). Meanwhile, polybrene (5µg/ml) was added to the cell culture medium. Retrovirus supernatant was then carefully removed from the plate and OT-I T cells were added to retrovirus-coated plate, spun for 10 min at 254 g and cultured with human IL-2, mouse IL-7 and mouse IL-15 (all at 5ng/ml) in complete RPMI for 2 days. For adoptive transfer experiments, 2 days after transduction GFP^+^ or mcherry^+^ cells were purified by flow cytometric cell sorting (BD FACS Aria^TM^ Fusion) prior to adoptive transfer into recipient mice. Of note, for experiments involving E2A overexpression, OT-I cells expressing a low to intermediate level of mCherry were FACS purified to avoid increased cell death in cells expressing higher level of E2A. Following T cell adoptive transfer recipient mice were then infected the following day with Tg.GRA6-OVA.

### mAb-depleting treatments

To deplete CD4^+^ T cells, an anti-CD4α mAb (rat IgG2b, clone GK1.5, cat # BX-BP0003 from BioXcell) or isotype (rat IgG2b, Clone LTF-2, cat # BP0090 from BioXcell) was injected intraperitoneally at day −3 and day −1 (100 μg per mouse) before infection and once weekly thereafter until the end of the experiment at day 35 post-*T.gondii* infection.

### Isolation of spleen and brain leukocytes

Mice received a lethal dose of ketamine (100 mg/kg) and xylazine (10 mg/kg) and then underwent intracardial perfusion with PBS 1X. Subsequently, spleen and brain were collected. Splenocytes were mashed through a 70 µm cell strainer (Falcon) in complete RPMI (GIBCO) supplemented with 10% FCS (GIBCO) and red blood cells were lysed in ACK. Brains were minced, homogenized, and digested for 45min in digestion solution composed of Hanks’ balanced salt solution supplemented with collagenase D (1 mg/ml, Roche Diagnostics cat # 54 11088882001) and deoxyribonuclease (DNase) I (20 μg/mL, Sigma-Aldrich cat # DN25). Cells were passed through a 70 µm cell strainer, which was washed with 5 ml of washing solution (Hank’s balanced solution containing 2% FCS and Hepes) and then cells were left to decant 12 minutes in 15 ml Falcon tube. After decantation 13ml out of the 15ml were collected and washed in 50 ml Hank’s washing solution and centrifuged 5 min at 607g. Cell pellet was then suspended in 30% Percoll (GE Healthcare) and centrifuged at 1590g for 30 minutes without brake. The pelleted cells were resuspended in FACS buffer (PBS 1X, 2% FCS), washed 2 times and utilized for subsequent experiments.

### Flow cytometry

To identify OVA-derived SIINFEKL peptide specifc-CD8^+^ T cells, splenocytes and brain leukocytes were incubated with PE-coupled SIINFEKL-loaded H-2 K^b^ dextramers (Immudex, dilution 1:50) in complete RPMI during 45 min at room temperature. Then, cells were stained with Fc receptor blocking antibodies (CD16/32, clone 93, dilution 1:5). Specific cell-surface staining was performed using a standard procedure with fluorochrome-labeled antibodies against CD8a, CD3e, CD45.1, CD45.2, Slamf6, Tim3, PD-1, CD69, CD103, CD49a, CD62L and CD44 (see details of antibodies in Supplementary Table 1). Cells were then labelled with eFluor 780 Fixable Viability Dye (eBioscience, dilution1/2000). For intracellular staining, Granzyme B, IFN-γ and TNF were detected using Cytofix/Cytoperm kit (BD Biosciences) while Tox staining were performed using Foxp3/Transcription Factor Staining Buffer Set (eBioscience) following the manufacturer’s protocol. Analysis was performed on a FACS LSR Fortessa (BD Biosciences) and data were analyzed using FlowJo flow cytometry analysis software version 10.9.0. For analysis of phenotypic markers, mice with less than 20 events in the relevant gate were excluded from the analysis.

### In vitro generation of exhausted CD8^+^ T cells

Protocol was adapted from (*41*). Briefly, CD8^+^ T cells were purified from spleens and inguinal LNs of OT-I mice by negative selection with magnetic beads. In each well of a 24-well plate, 5×10^5^ of the purified CD8^+^ T cells were cultured in complete RPMI medium supplemented with IL-15 (5ng/ml, Peprotech, Cat 210–15) and IL-7 (5ng/ml, Peprotech, Cat 210–07) with 10ng/ml OVA peptide. For single peptide stimulation, cells were cultured in the presence of OVA peptide for 48h hours. The peptide was then removed by washing the cells two times with complete medium on day 2. For the remaining 5 days, the cells were cultured in the complete medium with cytokines. For repeated peptide stimulation, 10ng/ml OVA peptide was added daily for 7 days. To ensure comparable culture conditions, the cells were washed on day 2, similar to the procedure used for single peptide stimulation. When the cells reached maximum density, they were split and cultured with fresh complete medium containing cytokines.

### T cell restimulation for intracellular cytokine assessment

For *ex vivo* stimulation one third of Percoll-isolated brain leukocytes were incubated for 4h at 37°C with Phorbol 12-Myristate 13-Acetate (PMA) (50 ng/ml) and ionomycin (1000 ng/ml) (Sigma-Aldrich) in presence of monensin (Golgistop) (dilution 1/1428 from eBioscience). For *in vitro* exhaustion experiments, 5.10^5^– 8.10^5^ OT-I CD8^+^ T cells were incubated for 4h at 37°C with 10µg/ml of OVA peptide in presence of monensin (Golgistop) (1/1428) (eBioscience).

### Parasite load analysis

Parasite DNA quantification by qPCR was conducted on genomic DNA extracted from 5% of every brain homogenate using the DNEasy Blood & Tissue Kit (QIAGEN). As previously outlined (*58*), amplification of a 529-bp repeat element in the *T. gondii* genome was achieved using the TOX 9 and TOX 11 primers. The quantity of parasite genomes per mg of tissue DNA was determined by comparing it with a standard curve, which was generated using a known quantity of Pru tachyzoites. The Primers sequences were:

TOX 9 : 5’ –AGGAGAGATATCAGGACTGTAG

TOX 11 : 5’-GCGTCGTCTCGTCTAGATCG

### Single cell RNA-seq libraries

scRNA-seq dataset is from one experiment in which C57BL/6 mice were chronically infected with Tg.GRA6-OVA and analyzed at day 154pi. CD8^+^ T cells were isolated from pooled spleens or brains of 4 infected mice. CD8^+^ T cells were purified using MACS Miltenyi mouse CD8α T Cell Isolation kit (Miltenyi Biotec 130-104-075) before proceeding to single-cell emulsion with 15946 splenic CD8^+^ T cells and 11856 brain CD8^+^ T cells, respectively. Single-cell libraries were generated with Chromium Next GEM Single Cell 5’ Kit v2 according to the manufacturer’s protocol (10X Genomics). Library size and quality were verified on a Fragment Analyzer (Agilent). Libraries were sequenced on 1 lane of SP flowcell with Illumina NovaSeq6000 sequencer, using paired-end sequencing (2 x 150) and a single 8 bp-long index.

### Bioinformatical analysis of single-cell RNA-seq data

#### Single-cell RNA sequencing data analysis

Single-cell RNA-seq data were sequenced and profiled using the Chromium 10X technology. Fastq files processing was carried out with CellRanger 10X software (version 4.0.0) (*59*) and reads were mapped against mouse reference genome (version mm10-2020-A). Filtered barcode matrix generated from CellRanger was then loaded into R (version 4.0.3) (R Core Team. R Foundation for Statistical Computing, 2022) and used as input for Seurat R package (version 4.2.1) x(*60*). Multiple filters were applied to keep cells of interest : first filter to exclude low-quality, dying cells and doublets given sample-specific thresholds on feature, UMI and mitochondrial transcript count per cell. Second filter aims to remove non lymphoid cells based on cell cluster marker genes, and cell type identification with clustifyR R library (version 1.10.0). Data were processed following the standard Seurat pipeline : gene were normalized and scaled with NormalizeData and ScaleData Seurat function, highly variable genes were selected with FindVariableFeatures function, we then reduced the dimensionality of our data by PCA and clustered our cells using FindNeighbors, FindClusters, RunUMAP Seurat functions. We identified differentially expressed genes for cluster and sample with FindAllMarkers and FindMarkers function. Functional enrichment analysis was performed with fgsea package (*61*) (v1.24.0) or the GSEA software (*62*) (http://www.broad.mit.edu/gsea/) with published gene signatures (*11, 25, 35*).

#### Statistical analyses

Statistical analyses for all experiments were performed with Prism software v10.1.2 (GraphPad). The normality of all datasets was evaluated using Shapiro test. If the data were normally distributed two-tailed unpaired t-test or paired t-tests were conducted. If the data were not normally distributed, non-parametric tests (Mann-Whitney test or Wilcoxon matched-pairs signed rank) were utilized. Tests were corrected for multiple comparisons when applicable. *P* values ≤0.05 were considered statistically significant and are indicated on the figures as follows: \**P* < 0.05, \*\**P* < 0.01, \*\*\**P* < 0.001, and \*\*\*\**P* < 0.0001.

## ACKNOWLEDGEMENTS

We are indebted to Dr Gabrielle T. Belz (The Frazer Institute, The University of Queensland, Brisbane, Australia) for the gift of the Id2 floxed mice as well as pMSCV-Id2-IRES-GFP, pMSCV-E47-IRES-mcherry and related control plasmids. We are grateful to Dr Klaas Van Gisbergen (Champalimaud Research, Champalimaud Foundation, Lisbon, Portugal) for the gift of the Hobit-Tdtomato-Cre-DTR reporter mice, Dr Corinne Mercier for the gift of the pGRA2/Ble/GRA2 plasmid, and Dr Sebastian Lourido for expert advice on the generation of ΔGRA2 parasites. Flow cytometry experiments were done at the INFINITy-INSERM UMR1291 core facility connected to Toulouse Réseau Imagerie network, member of the France-BioImaging national infrastructure supported by the French National Research Agency (ANR-10-INBS-04). We are very thankful to Hugo Garnier, Anne-Laure Iscache, Valerie Duplan and Fatima L’Faqihi for their assistance in cell sorting. We thank the UMS06 for the management of the animal facility and help with animal experimentation. We also thank the platform GetSanté (GENOTOUL Toulouse, France), in particular Frédéric Martins and Emeline Lhuillier for their help with the single-cell RNA sequencing libraries.

## CONTRIBUTIONS

### Author contributions

FM conceived the study. AC, with assistance from LN, CT, RP, EB, AH, and MB, carried out all the experiments. MA and NB analyzed the single-cell RNA-seq data. FM and AC were involved in data analysis. FM drafted the initial version of the paper, which was subsequently edited by AC and NB.

