## Supplementary data for "Id2 levels determine the development of effector vs. exhausted tissue-resident memory CD8^+^ T cells during CNS chronic infection"

### SUPPLEMENTARY MATERIAL

#### Supplementary Figures

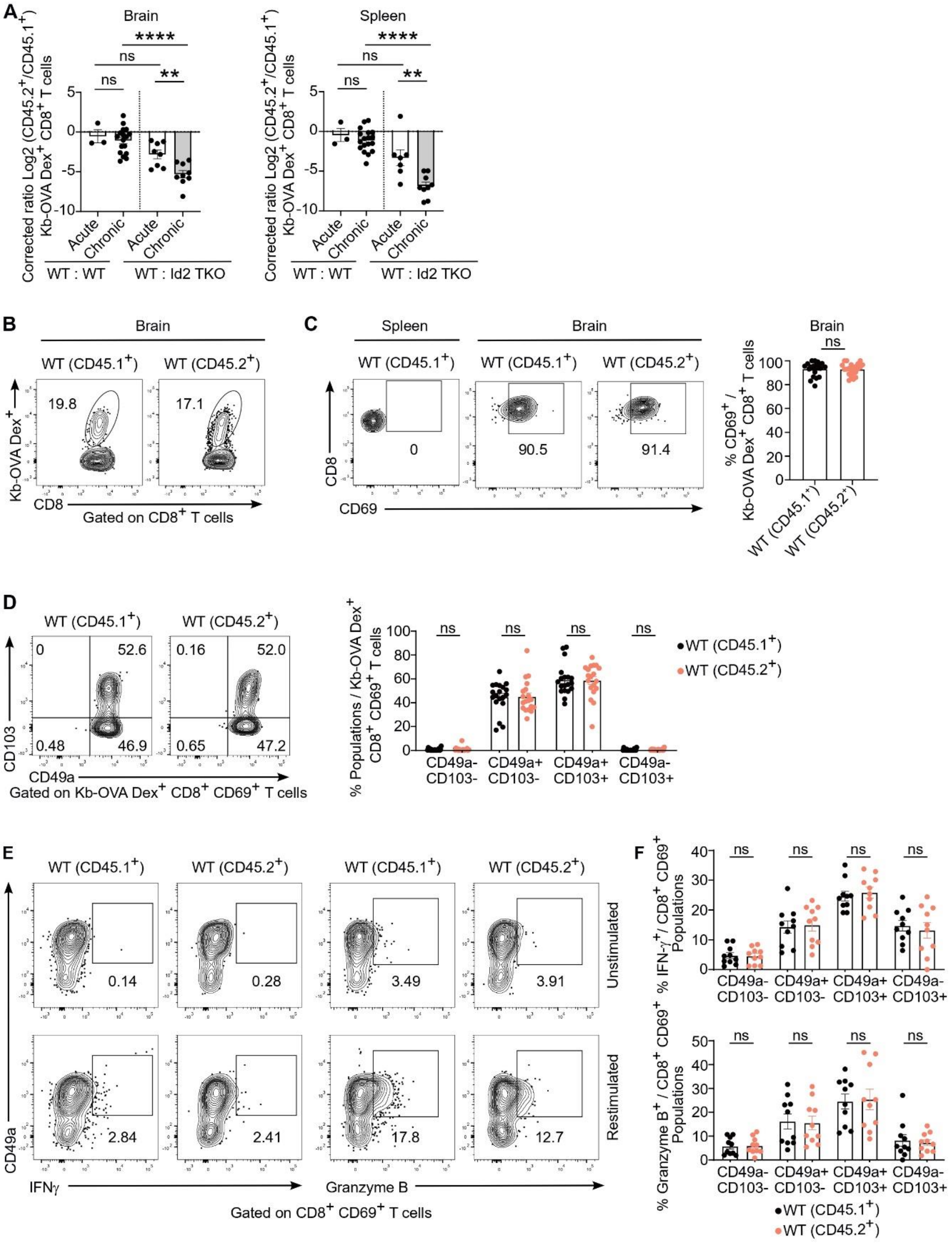

**Supplementary Figure 1. Control chimeras do not show defect in Trm development during the chronic phase of *T.gondii* infection.**

(**A**) Mixed bone marrow (BM) chimeras were reconstituted with a mix of Id2 TKO or WT ($CD45.2^+$) BM from Id2floxed;CD4Cre littermates and BM from WT ($CD45.1^+$) mice and were infected with Tg.ΔGRA2.GRA6-OVA 7-8 weeks post-BM injection. Mice were analyzed either 2 wks (acute phase) or 9-11 wks (chronic phase) post-infection (p.i). Bar graphs show the $log_2$ ratio of WT or Id2 TKO ($CD45.2^+$) to WT ($CD45.1^+$) $K^b$-OVA $Dex^+$ $CD8^+$ T cells corrected for the ratio of $CD45.2^+$ to $CD45.1^+$ within the splenic naïve $CD8^+$ T cell ($CD62L^+$ $CD44^-$ $CD8^+$) compartment in the indicated organs of mixed bone marrow chimeras at acute and chronic stages. **(B-F**) Analysis of control mixed BM chimeras WT ($CD45.2^+$) : WT ($CD45.1^+$) 9-11 weeks post-infection with Tg.ΔGRA2.GRA6-OVA. (**B**) Representative contour plots show the frequency of WT ($CD45.2^+$) and WT ($CD45.1^+$) OVA-specific ($K^b$-OVA $Dex^+$) $CD8^+$ T cells within the brain. (**C**) Representative contour plots show CD69 expression within brain $K^b$-OVA $Dex^+$ $CD8^+$ T cells. Bar graph shows the percentage of $CD69^+$ cells among brain WT ($CD45.1^+$) and WT ($CD45.2^+$) OVA-specific $CD8^+$ T cells. (**D**) (Left) Representative contour plots of CD49a and CD103 expression gated on brain WT ($CD45.2^+$) or WT ($CD45.1^+$) $K^b$-OVA $Dex^+$ $CD69^+$ $CD8^+$ T cells. (Right) Bar graph shows the indicated population among brain WT ($CD45.2^+$) and WT ($CD45.1^+$) $CD69^+$ $K^b$-OVA $Dex^+$ $CD8^+$ T cells. (**E**) Representative contour plots show the intracellular expression of IFN-γ and granzyme B within brain WT ($CD45.2^+$) and WT ($CD45.1^+$) $CD44^+$ $CD69^{+.}$ $CD8^+$ T cells after restimulation either with PMA and ionomycin in presence of monensin, or with monensin alone. (**F**) Bar graphs show the frequency of IFN-γ (Top) and Granzyme B (Bottom) expressing cells after PMA/ionomycin restimulation among brain infiltrating total WT ($CD45.2^+$) or WT ($CD45.1^+$) $CD8^+$ $CD69^+$ T cells. Data in **A** are pooled from 3 experiments (acute phase) to 4 experiments (chronic phase) for Id2 TKO:WT chimeras, and from 1 experiment (acute phase) to 4 experiments (chronic phase) for WT: WT control chimeras. Data in **C-F** are pooled from 2 (E, F) to 4 (C-D) independent experiments. Bars show the mean ± SEM. Statistically significant differences were determined using one-way ANOVA (A), Wilcoxon matched-pairs signed rank tests (C) and multiple Wilcoxon matched-pairs signed rank testys corrected for multiple comparisons using Hom-Sidak method (D, F). **P* < 0.05, ***P* < 0.01, ****P* < 0.001, and *****P* < 0.0001. ns, not significant. Each dot represents an individual mouse.

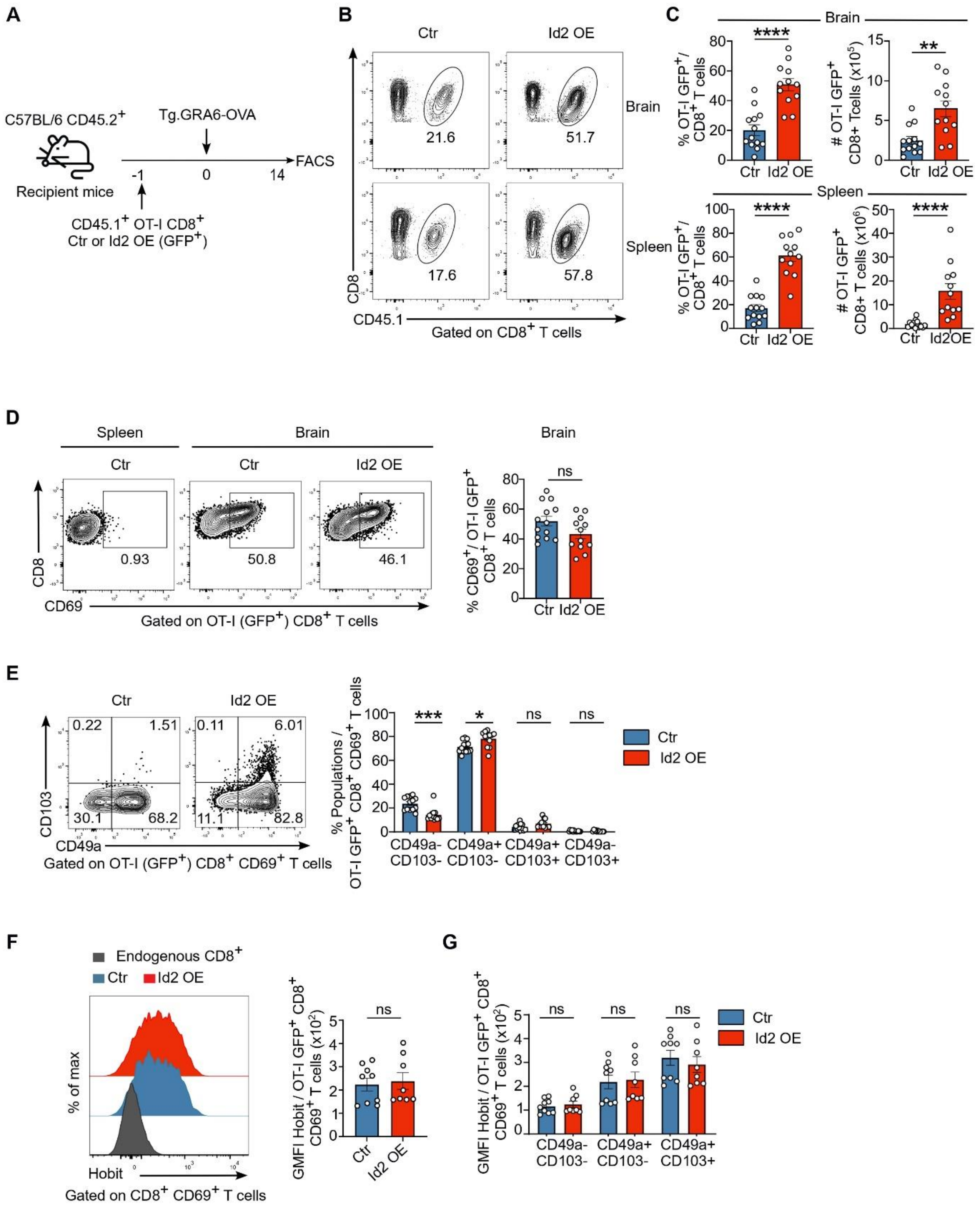

**Supplementary Figure 2 : Id2 overexpression increases the formation of CD49a$^+$ brain Trm during the acute phase of *T.gondii* infection**

(**A**) Schematic diagram outlining the adoptive transfer of TCR transgenic OVA-specific CD8$^+$ T cells (OT-I) transduced with retroviruses encoding Id2 (Id2 OE) or a control (Ctr) empty vector and the GFP as fluorescent reporter, to recipient mice subsequently infected with Tg.GRA6-OVA. Mice were analysed at day 14 p.i and brains were collected. (**B**) Representative contour plots show the percentage of Ctr and Id2 OE OT-I GFP$^+$ among total CD8$^+$ T cells from the brain (top) and spleen (bottom). (**C**) Bar graphs show the frequency among CD8$^+$ T cells (left) and total number (right) of Ctr and Id2 OE OT-I GFP$^+$ (CD45.1$^+$) in the brain and spleen. (**D**) Representative contour plots (left) and bar graph (right) show the percentage of CD69$^+$ cells among brain Ctr and Id2 OE OT-I GFP$^+$ (CD45.1$^+$). (**E**) (Left) Representative contour plots show the expression of CD103 vs. CD49a in brain Ctr and Id2 OE CD69$^+$ OT-I GFP$^+$ (CD45.1$^+$) CD8$^+$ T cells. (Right) Bar graph shows the percentage of the indicated population within brain Ctr and Id2 OE CD69$^+$ OT-I GFP$^+$ (CD45.1$^+$) cells. **(F)** (Left) Representative flow cytometric histogram shows the expression of Hobit-tdTomato in brain Ctr (blue) and Id2 OE (red) GFP$^+$ CD69$^+$ OT-I x Hobit-tdTomato (CD45.1$^+$) CD8$^+$ T cells and endogenous CD8$^+$ T cells (black). Bar graph shows the GMFI of Hobit-tdTomato of brain Ctr and Id2 OE GFP$^+$ CD69$^+$ OT-I x Hobit-tdTomato (CD45.1$^+$) cells. **(G)** Bar graph shows the GMFI of Hobit-tdTomato in the indicated population within brain Ctr and Id2 OE GFP$^+$ CD69$^+$ OT-I x Hobit-tdTomato (CD45.1$^+$) cells. Data are pooled from 2 (F, G) and 3 (C, D, E) independent experiments. Bars show the mean ± SEM. Statistically significant differences were determined using Mann-Whitney (C, D, F) and multiple Mann-Whitney (E, G) tests corrected for multiple comparisons using Hom-Sidak method. *$P < 0.05$, **$P < 0.01$, ***$P < 0.001$, and ****$P < 0.0001$. ns, not significant. Each dot represents an individual mouse.

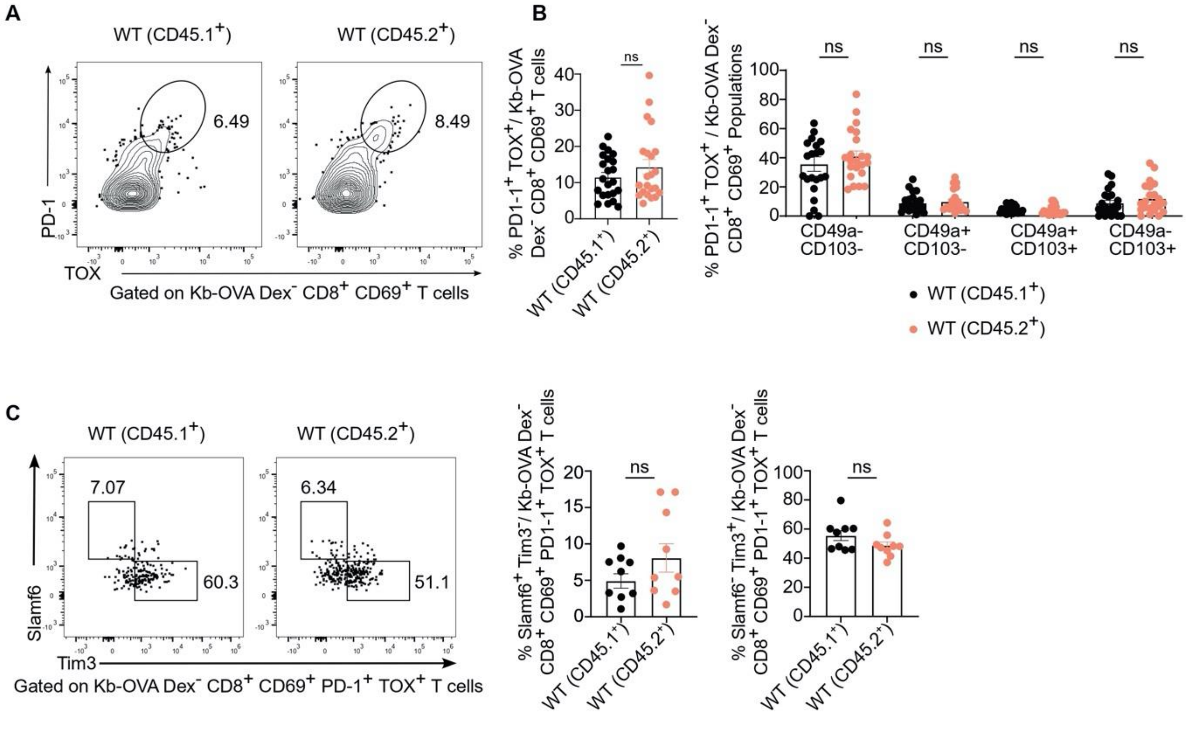

**Supplementary Figure 3 : No difference of T cell exhaustion in control mixed chimeras during the chronic phase of *T.gondii* infection.**

Control mixed BM chimeras WT (CD45.2$^+$):WT (CD45.1$^+$) were generated as described in Fig. 2A and were analyzed during the chronic phase (9-11 wks p.i.) of Tg.ΔGRA2.GRA6-OVA infection. **(A)** Representative contour plots of PD-1 and Tox expression in WT (CD45.1$^+$) and WT (CD45.2$^+$) total CD8$^+$ CD44$^+$ T cells from the brain during the chronic phase of the infection. (**B**) (left) Bar graph shows the frequency of PD-1$^+$TOX$^+$ expressing cells among WT (CD45.1$^+$) and WT (CD45.2$^+$) CD8$^+$ CD44$^+$ CD69$^+$ T cells from the brain at the chronic stage of the infection. (right) Bar graph shows the percentage of PD-1$^+$TOX$^+$ expressing cells within the indicated populations among WT (CD45.1$^+$) and WT (CD45.2$^+$) CD69$^+$ CD44$^+$ CD8$^+$ T cells in the brain during the chronic phase of the infection. (**C**) Representative dot plots (left) and bar graphs (right) show the proportion of Slamf6$^+$Tim3$^-$ and Slamf6$^-$Tim3$^+$ cells among brain WT (CD45.2$^+$) and WT (CD45.1$^+$) PD-1$^+$ Tox$^+$ CD69$^+$ CD44$^+$ CD8$^+$ T cells. Data are pooled from 2 (C), and 5 (B) independent experiments. Bars show the mean ± SEM. Statistically significant differences were determined using Paired t-test (C), Wilcoxon matched-pairs signed rank tests (B left) and multiple Paired matched-pairs signed rank tests corrected for multiple comparisons using Hom-Sidak method (B right). *$P < 0.05$, **$P < 0.01$, ***$P < 0.005$. ns, not significant. Each dot represents an individual mouse.

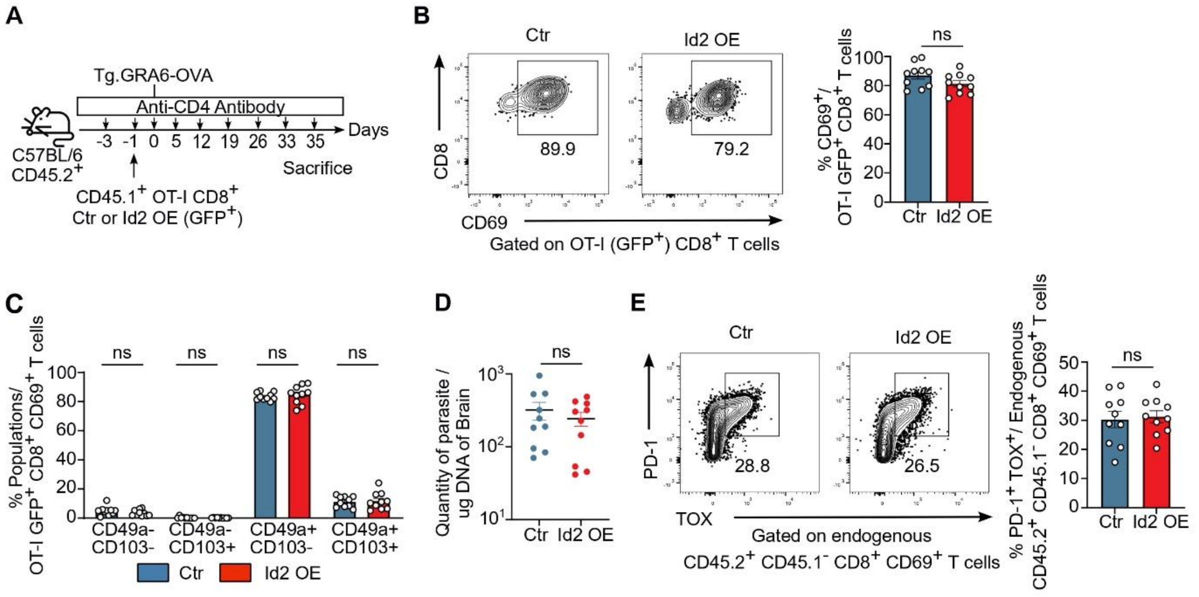

**Supplementary Figure 4 : No differences in Trm adhesion molecules expression and parasite load in mice transferred with Id2 OE and Ctr OT-I cells upon CD4 T cell depletion during the chronic phase of *T.gondii* infection.**

(**A**) Mice were adoptively transferred with OT-I (CD45.1$^+$) cells transduced with a control vector or an Id2 overexpressing vector expressing the GFP reporter one day prior Tg.GRA6-OVA infection. All mice were treated with the anti-CD4 depleting antibody at day -3 and -1 before infection and CD4$^+$ T cell depletion was repeated once per week until day 35 p.i. (**B**) Representative contour plots (left) and bar graph (right) show the percentage of CD69$^+$ cells among brain Ctr and Id2 OE OT-I GFP$^+$ (CD45.1$^+$) CD8$^+$ T cells (**C**) Bar graph shows the frequency of the indicated population among brain Id2 OE and Ctr CD69$^+$ OT-I GFP$^+$ (CD45.1$^+$). (**D**) Bar graph shows parasite burden measured by qPCR on genomic DNA extracted from the brains of mice transferred with Ctr or Id2 OE OT-I GFP$^+$ (CD45.1$^+$) CD8$^+$ T cells. Representative flow cytometric histogram and bar graph show PD-1 expression among total CD69$^+$ brain infiltrating OT-I (CD45.1$^+$). (**E**) Representative contour plots (left) and bar graph (right) show the percentage of PD1$^+$ Tox$^+$ cells among brain endogenous CD8$^+$ T cells

($CD44^{+}$ $CD45.1^{-}$) from mice transferred with Ctr and Id2 OE OT-I $GFP^{+}$ ($CD45.1^{+}$). Data are pooled from 2 (B, C, D, E) independent experiments. Bars show the mean ± SEM. Statistically significant differences were determined using Mann-whitney (B, D, E) and multiple Mann-whitney tests corrected for multiple comparisons using Hom-Sidak method (C). *$P < 0.05$, **$P < 0.01$, ***$P < 0.001$, and ****$P < 0.0001$. ns, not significant. Each dot represents an individual mouse.

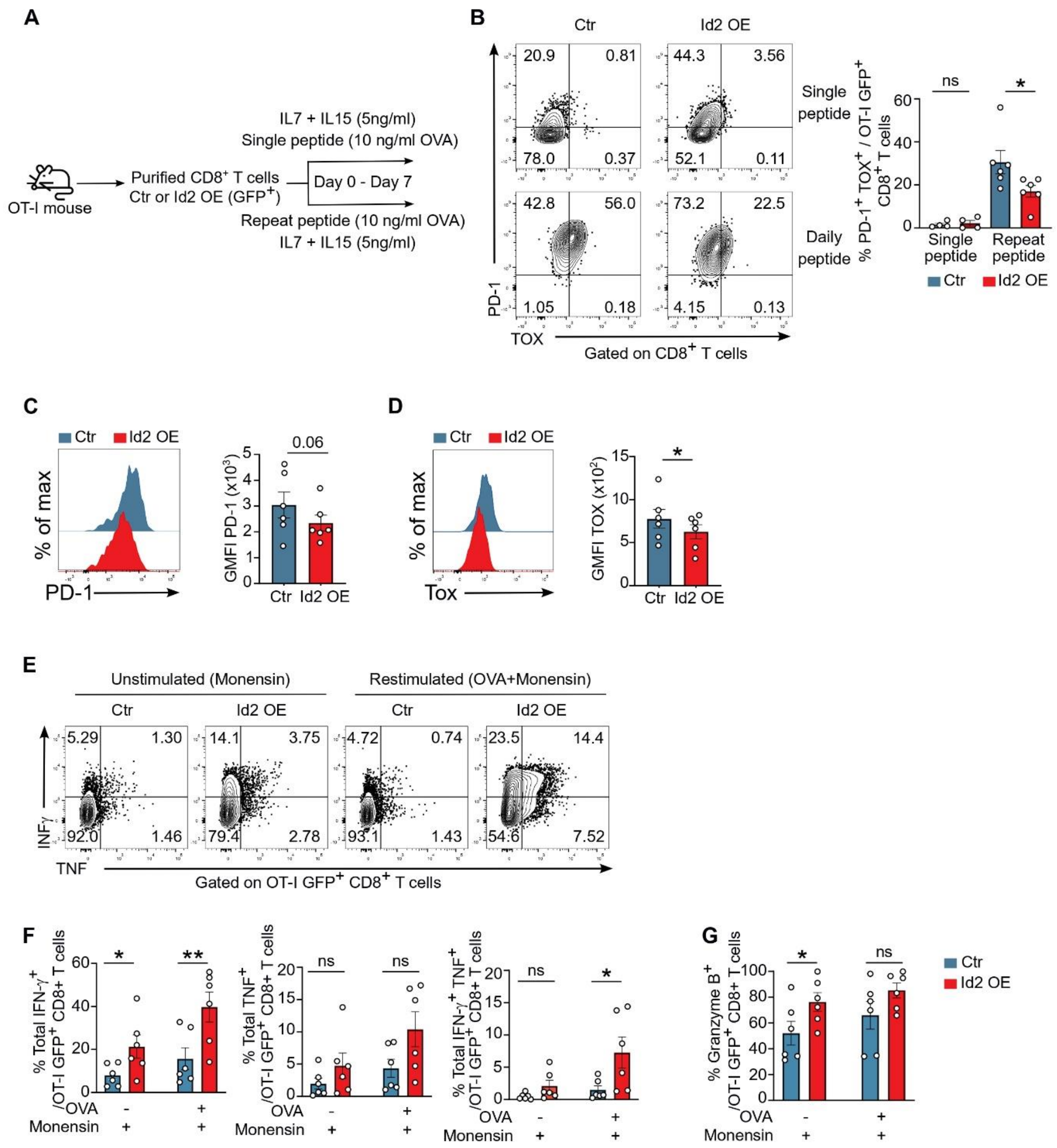

**Supplementary Figure 5 : Id2 overexpression decreases T cell exhaustion in a model of *in vitro* induced T cell exhaustion.**

(**A**) Experimental workflow diagram : Naive Tcr transgenic OVA-specific CD8$^+$ T cells (OT-I) were isolated from the spleen and lymph nodes, then transduced with either Id2-IRES-GFP (Id2 OE) or with Empty-IRES-GFP vector (Ctr). OT-I cells were stimulated once (Single peptide) or everyday (Repeat peptide) in presence of IL-7 and IL-15. At day 7 after activation GFP$^+$ cells were FACS-sorted. (**B**) Representative contour plots and bar graph show the proportion of PD-1$^+$TOX$^+$ expressing cells at day 7 in FACS-sorted GFP$^+$ Ctr and Id2 OE OT-I cells. (**C**) Representative flow cytometric histogram and bar graph show PD-1 GFMFI among Ctr and Id2 OE OT-I. (**D**) Representative flow cytometric histogram and bar graph show Tox GFMI among Ctr and Id2 OE OT-I. (**E**) Representative contour plots show intracellular IFN-γ and TNF expression after restimulation with OVA peptide + monensin or with monensin alone. (**F**) Bar graphs show the percentage of cells expressing IFN-γ or TNF or co-expressing IFN-γ and TNF cells after OVA peptide restimulation. (**G**) Bar graphs show the percentage granzyme B expressing cells after OVA peptide restimulation. Data are pooled from 3 (B-G) independent experiments. Bars show the mean ± SEM. Statically significant differences were determined using Paired t-tests (B, C, D, F, G). *$P < 0.05$, **$P < 0.01$, ***$P < 0.001$, and ****$P < 0.0001$. ns, not significant. Each dot represents an individual mouse.

**Supplementary Table 1** : Antibodies used for flow cytometric analysis

| **Antibodies** | **Fluorochrome** | **Dilution** | **Souce** | **Identifier** |
|---|---|---|---|---|
| CD45.1 | EF450 | 1/200 | Thermofisher | Cat # 48-0453-82 Clone A20 |
| CD45.1 | Alexa Fluor® 700 | 1/200 | BD | Cat # 561235 Clone A20 |
| CD45.1 | BUV496 | 1/200 | BD | Cat # 741093 Clone A20 |
| CD45.2 | PerCP-Cy 5.5 | 1/200 | BD | Cat # 552950 Clone 104 |
| CD45.2 | Alexa Fluor® 700 | 1/200 | BD | Cat # 560693 Clone 104 |
| CD45.2 | Alexa Fluor® 488 | 1/200 | Biolegend | Cat # 109816 Clone 104 |
| CD8 | BV786 | 1/300 | BD | Cat # 563332 Clone 53-6.7 |
| CD8 | BV421 | 1/300 | BD | Cat # 563898 Clone 53-6,7 |
| CD8 | APC | 1/300 | BD | Cat # 553035 Clone 53-6,7 |
| CD8 | PE-Cy7 | 1/300 | Biolegend | Cat # 100722 Clone 53-6,7 |
| CD8 | BV805 | 1/500 | BD | Cat # 612898 Clone 53-6,7 |
| CD3e | Alexa Fluor® 700 | 1/300 | BD | Cat # 561388 Clone 17A2 |
| CD3e | BV711 | 1/200 | Biolegend | Cat # 100349 Clone 145-2C11 |
| CD62L | APC | 1/400 | BD | Cat # 553152 Clone MEL-14 |

| | | | | |
|---|---|---|---|---|
| CD62L | BUV563 | 1/500 | BD | Cat 741230 Clone MEL-14 |
| CD44 | Alexa Fluor® 700 | 1/400 | Thermofisher | Cat # 56-0441-82 Clone IM7 |
| CD44 | PerCP-Cy 5.5 | 1/400 | Thermofisher | Cat # 45-0441-82 Clone IM7 |
| CD44 | BV570 | 1/200 | Biolegend | Cat # 103037 Clone IM7 |
| PD-1 (CD279) | PE-Dazzle 594 | 1/200 | Biolegend | Cat # 109116 Clone RMP1-30 |
| PD-1 (CD279) | BV786 | 1/200 | BD | Cat # 744548 Clone J43 |
| Slamf6 (Ly108) | BV786 | 1/200 | BD | Cat # 741030 Clone 13G3 |
| Slamf6 (Ly108) | Pacific blue | 1/200 | Biolegend | Cat # 134608 Clone 330-AJ |
| Tim3 (CD366) | BB700 | 1/200 | BD | Cat # 747619 Clone 5D12/TIM-3 |
| CD69 | PE-Cy7 | 1/200 | Thermofisher | Cat # 25-0691-82 Clone H1.2F3 |
| CD69 | BUV615 | 1/100 | BD | Cat # 751593 Clone H1.2F3 |
| CD103 | BV510 | 1/200 | BD | Cat # 563087 Clone M290 |
| CD49a | BUV737 | 1/400 | BD | Cat # 741776 Clone Ha31/8 |
| TOX | APC | 1/100 | Miltenyi | Cat # 130-118-335 Clone REA473 |

| | | | | |
|---|---|---|---|---|
| Granzyme B | PE | 1/200 | Thermofisher | Cat # 12-8898-82 Clone NGZB |
| Granzyme B | Alexa Fluor® 647 | 1/200 | Biolegend | Cat # 515406 Clone GB11 |
| IFN-g | BV786 | 1/200 | BD | Cat # 563773 Clone XMG1.2 |
| TNF | Alexa Fluor® 700 | 1/200 | BD | Cat # 558000 Clone MP6-XT22 |
